# Ether lipids influence cancer cell fate by modulating iron uptake

**DOI:** 10.1101/2024.03.20.585922

**Authors:** Ryan P. Mansell, Sebastian Müller, Jia-Shu Yang, Sarah Innes-Gold, Sunny Das, Ferenc Reinhardt, Kim Janina Sigmund, Vaishnavi V. Phadnis, Zhengpeng Wan, Jillian Stark, Daehee Hwang, Fabien Sindikubwabo, Laurène Syx, Nicolas Servant, Elinor Ng Eaton, Julio L. Sampaio, George W. Bell, Prathapan Thiru, Amartya Viravalli, Amy Deik, Clary B. Clish, Paula T. Hammond, Roger D. Kamm, Adam E. Cohen, Ilya Levental, Natalie Boehnke, Victor W. Hsu, Kandice R. Levental, Raphaël Rodriguez, Robert A. Weinberg, Whitney S. Henry

## Abstract

Cancer cell fate has been widely ascribed to mutational changes within proteincoding genes associated with tumor suppressors and oncogenes. In contrast, the mechanisms through which the biophysical properties of membrane lipids influence cancer cell survival, dedifferentiation and metastasis have received little scrutiny. Here, we report that cancer cells endowed with high metastatic ability and cancer stem cell-like traits employ ether lipids to maintain low membrane tension and high membrane fluidity. Using genetic approaches and lipid reconstitution assays, we show that these ether lipid-regulated biophysical properties permit non-clathrin-mediated iron endocytosis via CD44, resulting in significant increases in intracellular redox-active iron and enhanced ferroptosis susceptibility. Using a combination of in vitro threedimensional microvascular network systems and in vivo animal models, we show that loss of ether lipids from plasma membranes also strongly attenuates extravasation, metastatic burden and cancer stemness. These findings illuminate a mechanism whereby ether lipids in carcinoma cells serve as key regulators of malignant progression while conferring a unique vulnerability that can be exploited for therapeutic intervention.

## INTRODUCTION

Cancer cells have the capacity to undergo dynamic changes in identity, structure, and function, making them remarkably versatile and adaptable. Alterations in the lipid composition of cell membranes contribute to the phenotypic plasticity of cells. The distinctive physicochemical properties and subcellular localization of various lipids within cell membranes influence a range of biological processes, including cellular trafficking, signaling and metabolism^1^. Despite growing knowledge of lipid biology, an understanding of how specific lipid subtypes impact cancer cell fate remains limited.

Emerging studies demonstrate that therapy-resistant mesenchymal-like carcinoma cells exhibit an elevated vulnerability to ferroptosis^2–4^, an iron-dependent form of cell death characterized by an unrestricted accumulation of oxidized membrane phospholipids^5–8^. In previous work we showed that the natural product salinomycin can selectively eliminate otherwise therapy-resistant, mesenchymal-enriched cancer stem cells (CSC) by targeting lysosomal iron to promote an iron-dependent cell death^4, 9, 10^. In this context, we found that such CSC-enriched cells exhibit a high intracellular iron load compared to their non-CSC-like counterparts, rendering them especially vulnerable to elimination by induced ferroptosis^4^.

Ferroptosis can also be instigated by pharmacologic inhibition of ferroptosis suppressors, such as glutathione peroxidase 4 (GPX4)^5, 11^, ferroptosis-suppressor protein 1 (FSP1, previously known as AIFM2)^12, 13^, as well as through downregulation of reduced glutathione (GSH)^14, 15^. Activated CD8^+^ T cells may also induce ferroptosis in cancer cells^16, 17^. Beyond cancer, ferroptosis has been implicated in the pathogenesis of several neurodegenerative diseases and acute injury of the kidney, liver and heart^18–21^.

In previous work, we undertook an unbiased, genome-wide CRISPR/Cas9 screen with the goal of identifying genes that govern ferroptosis susceptibility in highgrade human serous ovarian cancer cells^22^. This screen revealed a previously unrecognized role for ether lipid-synthesizing enzymes, such as alkylglycerone phosphate synthase (AGPS), in modulating ferroptosis susceptibility. The ether phospholipids generated by these enzymes represent a unique subclass of glycerophospholipids characterized by an ether-linked hydrocarbon group formed at the *sn*-1 position of the glycerol backbone^23^. This phospholipid subtype constitutes ∼20% of the total phospholipid pool in many types of mammalian cells.

The significance of ether lipid species in human health is underscored by the severe inherited peroxisomal disorders caused by their deficiency^23^. This often manifests as profound developmental abnormalities, such as neurological defects, vision and hearing loss, and reduced lifespan. In the context of cancer, elevated ether lipid levels have been correlated with increased metastatic potential of carcinoma cells^24–27^. Despite these pathological associations, the mechanism(s) by which ether lipids affect cancer progression remain(s) elusive. Furthermore, the mechanism(s) by which loss of ether lipids results in decreased ferroptosis susceptibility required further investigation.

Our previous work, along with that of others, ascribed a role to polyunsaturated ether phospholipids as chemical substrates prone to the iron-mediated oxidation that triggers ferroptotic cell death^22, 28^. Here, we demonstrate that ether lipids also play an unrelated biophysical role, facilitating iron endocytosis in carcinoma cells. This represents an unappreciated mechanism of intracellular signaling by which ether lipids contribute to intracellular levels of a critical metal catalyst. In addition, our findings highlight the functional importance of this poorly studied lipid subtype in enabling a variety of malignancy-associated cell phenotypes, including metastasis and tumor-initiating abilities. Taken together, these results establish a role for ether lipids as critical effectors of cancer cell fate.

## RESULTS

### Ether lipids play a key role in maintaining a ferroptosis-susceptible cell state

To investigate the mechanism(s) by which ether lipid deficiency reduces ferroptosis susceptibility, we employed CRISPR/Cas9 to knockout (KO) the *AGPS* gene in ferroptosis-sensitive TGF-β-treated PyMT-1099 murine breast carcinoma cancer cells^29^ (Supplementary Fig. 1a). The *AGPS* gene encodes a rate-limiting enzyme critical for ether lipid biosynthesis^23^ (Fig. 1a). Consistent with our prior studies^22^, loss of ether lipids via AGPS KO significantly decreased the susceptibility of these cancer cells to ferroptosis induced by treatment with the GPX4 inhibitors RSL3 or ML210 (Fig. 1b, Supplementary Fig. 1b).

**Fig. 1:**
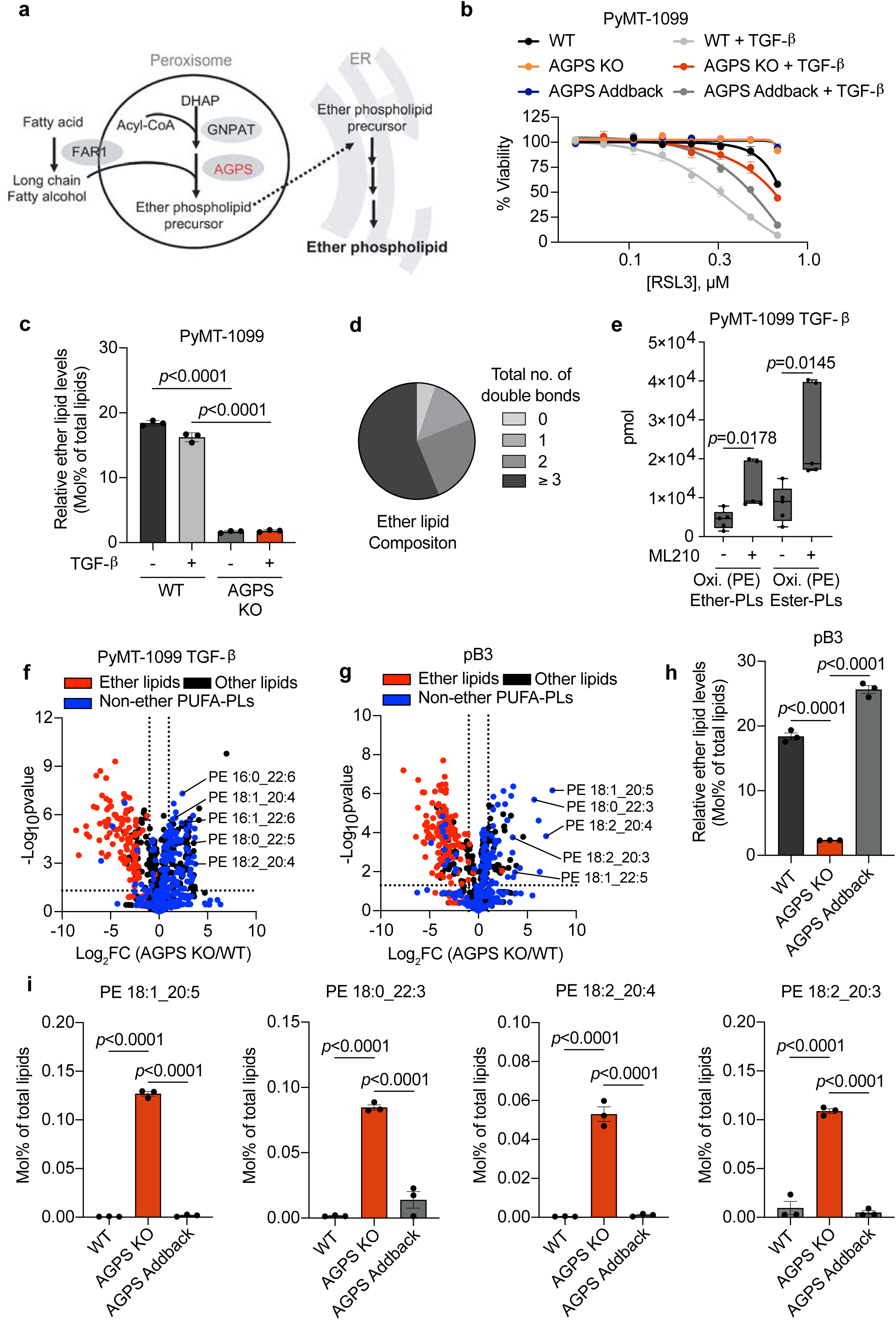
Ether lipids play a key role in maintaining a ferroptosis susceptible cell state. a. Schematic of peroxisomal-ether lipid biosynthetic pathway. b. Cell viability following treatment with the GPX4 inhibitor RSL3 for 72 h. PyMT-1099 WT, AGPS KO, or AGPS addback cells were pretreated with TGF-β (2 ng/ml) for 10 d prior to assay. Data shown as mean +/− SEM. Graph is representative of three independent biological replicates. c. Bar graph showing percent of total lipids constituted by ether lipids following AGPS KO in untreated wildtype (WT) or TGF-β-treated (2 ng/ml;10 d) PyMT-1099 cells. d. Pie chart showing the relative proportion of ether lipids with various total numbers of double bonds. e. Amount in pmol of oxidized phosphatidylethanolamine (Oxi. PE) ether and ester phospholipids in PyMT-1099 TGF-β cells treated with ML210 or vehicle control for 24 h. Five biological replicates per condition. 100,000 cells were used for lipid extraction in each condition. Statistical significance was calculated by unpaired, twotailed t-test; ns, not significant. f. Volcano plot showing the log_2_ fold change in the relative abundance of various lipid species upon knockout of AGPS in PyMT-1099 TGF-β-treated cells. Blue indicates non-ether linked polyunsaturated phospholipids (PUFA-PLs), red indicates all ether lipids identified in lipidomic analysis and black denotes all other lipids identified. g. Volcano plot showing the log_2_ fold change in the relative abundance of various lipid species upon knockout of AGPS in pB3 cells. Blue indicates non-ether linked polyunsaturated phospholipids (PUFA-PLs), red indicates all ether lipids identified in lipidomic analysis and black denotes all other lipids identified. h. Bar graph showing the percent of total lipids constituted by ether lipids in pB3 WT, pB3 AGPS KO and pB3 AGPS addback cells. i. Bar graphs showing the effects of ether lipids on the relative abundance of selected polyunsaturated diacyl phospholipids in pB3 cells. All lipidomic data were analyzed in triplicate and shown as the mean +/− SEM. Unless otherwise stated statistical significance was calculated by one-way ANOVA with Tukey’s multiple comparisons test; ns, not significant. For panel b: PyMT-1099 WT and AGPS KO cells were transduced with the respective vector control plasmids. For panels h-i: pB3 WT and AGPS KO cells were transduced with the respective vector control plasmids.

Lipidomic analysis confirmed that *AGPS* knockout significantly reduces total ether lipid abundance, while only marginally affecting most other membrane-associated lipid species (Fig. 1c, Supplementary Fig. 2a). More than half of the identified ether lipids contained polyunsaturated fatty acyl groups, which are highly prone to hydrogen abstraction by reactive free radicals (Fig. 1d). Based on this observation, we speculated that loss of ether lipids may attenuate ferroptosis susceptibility by depleting the pool of available ether lipid substrates for lipid peroxidation. Thus, we performed oxidized lipidomic analysis on two ferroptosis-sensitive breast cancer cell lines treated with either ML210 or RSL3. These experiments confirmed that ether lipids are oxidized following ferroptosis induction (Fig. 1e, Supplementary Fig. 1c).

In order to test whether ether lipid deficiency leads to a global reduction of cellular pools of oxidation prone lipid species, we also investigated whether the relative abundance of pro-ferroptosis non-ether-linked polyunsaturated phospholipids was impacted by loss of ether lipids. Our analyses revealed that while total levels of phospholipid unsaturation are only marginally changed, ether lipid deficiency increased the relative abundance of several polyunsaturated diacyl phosphoethanolamine (PE) and phosphatidylcholine (PC)-derived lipids with putative pro-ferroptosis function^30, 31^ (Fig. 1f, Supplementary Fig. 2b-d). To ensure that these findings were not an idiosyncrasy of our TGF-β-treated PyMT-1099 AGPS KO cells, we confirmed this observation in PyMT-MMTV-derived pB3 murine AGPS KO breast carcinoma cells^32^ (Fig. 1g, 1h, Supplementary Fig. 1d, 3a-3f). Importantly, re-expression of AGPS (addback) restored the relative levels of these non-ether-linked polyunsaturated diacyl phospholipids to levels comparable to pB3 wildtype (WT) cells (Fig. 1i). These findings dispel the notion that ether lipid deficiency attenuates ferroptosis susceptibility simply by decreasing the global level of polyunsaturated phospholipids, further underscoring the importance of polyunsaturated ether phospholipids in maintaining a ferroptosis susceptible cell state.

### Ether lipids regulate levels of cellular redox-active iron in cancer cells

The above observations, together with our oxidized ether phospholipidomic analysis, supported the notion that ether lipids could modulate ferroptosis susceptibility— at least in part by serving as substrates for lipid peroxidation. However, these observations failed to address the possibility that alterations in ether lipid composition could also affect intracellular levels of redox-active iron, the central mediator of the lipid peroxidation that drives ferroptosis. Therefore, we used two orthogonal analyses to assess intracellular iron levels. Since the endolysosomal compartment is a key reservoir of reactive iron within cells^7, 33–35^, we used a lysosomal iron (II)-specific fluorescent probe, HRhoNox-M^36^, to gauge levels of lysosomal redox active iron. In addition, we used inductively coupled plasma mass spectrometry (ICPMS) to quantify total intracellular iron levels^7, 35, 37–39^.

Unexpectedly, loss of ether lipids (via AGPS KO) reduced intracellular iron levels in all murine cancer cell lines tested, whereas AGPS addback restored intracellular iron to levels comparable to those seen in parental ferroptosis-sensitive cancer cells (Fig. 2a-2d, Supplementary Fig. 4a). These findings provided the first indication that changes in ether phospholipids directly affect the levels of intracellular iron. Indeed, there was no direct precedent for the ability of a membrane-associated phospholipid to serve as a key regulator of the levels of an intracellular metal ion.

**Fig. 2:**
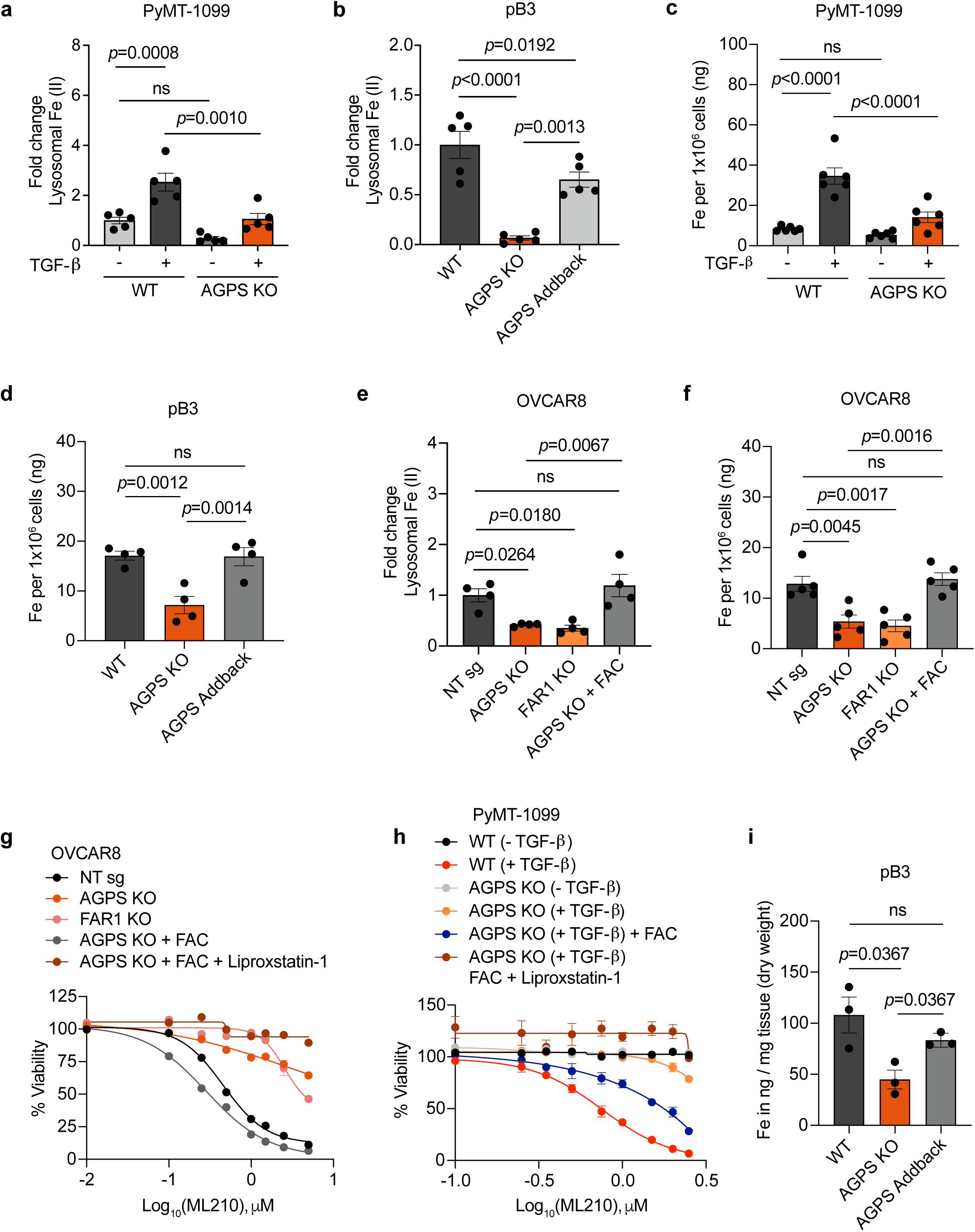
Ether lipids regulate cellular redox-active iron levels in cancer cells. a. Relative lysosomal iron levels (see methods) of PyMT-1099 WT or AGPS KO cells (+/−) pretreatment with TGF-β (n=5). Fold change calculated relative to the average of untreated PyMT-1099 wild-type (WT) cells. Statistical significance was calculated by one-way ANOVA with Holm-Šídák multiple comparisons test. b. Relative lysosomal iron levels of WT, AGPS KO, and AGPS Addback pB3 cells (n=5). Fold change calculated relative to the average of pB3 WT cells. Statistical significance was calculated by one-way ANOVA with Holm-Šídák multiple comparisons test. c. Inductively coupled plasma-mass spectrometry (ICP-MS) of cellular iron in PyMT-1099 WT or AGPS KO cells (+/−) pretreatment with TGF-β (n=6). Statistical significance was calculated by one-way ANOVA with Tukey’s multiple comparisons test. d. ICP-MS of cellular iron in WT, AGPS KO, and AGPS Addback pB3 cells (n=4). Statistical significance was calculated by one-way ANOVA with Tukey’s multiple comparisons test. e. Relative lysosomal iron levels in OVCAR8 NT sg, FAR1 KO or AGPS KO cells pretreated with ferric ammonium citrate (FAC, 50 μg/mL) (n=4). Fold change is calculated relative to the average of NT sg cells. Statistical significance was calculated by one-way ANOVA with Holm-Šídák multiple comparisons test. f. ICP-MS of cellular iron in OVCAR8 NT sg, FAR1 KO or AGPS KO cells pretreated with FAC (50 μg/mL) (n=5). Statistical significance was calculated by one-way ANOVA with Tukey’s multiple comparisons test. g. Cell viability of OVCAR8 NT sg, FAR1 KO or AGPS KO cells (+/−) FAC pretreatment followed by ML210 treatment for 72 h (n=3 technical replicates). Liproxstatin-1 (0.2 μM) was added at the time of ML210. Data representative of 3 independent experiments. h. Cell viability in response to ML210 treatment of PyMT-1099 WT or AGPS KO (+/−) pretreatment with TGF-β (10 d) followed by FAC treatment (100 µg/ml) for an additional 24 h. Cells were then treated with ML210 (+/−) liproxstatin-1 (0.2µM) and cell viability was assessed after 72 h (n=3 technical replicates). Data representative of 3 independent experiments. i. ICP-MS of cellular iron from primary tumors derived from WT, AGPS KO, and AGPS addback pB3 cells. Mean +/− SEM from 3 independent tumor samples per condition. Group differences were tested by permutation ANOVA followed by one tailed Welch’s t-tests with Holm correction for multiple comparisons. For panels b, d & i: pB3 WT and AGPS KO cells were transduced with the respective vector control plasmids. Abbreviation: NT – nontargeting, ns – not significant.

Supporting these findings, we found that the depletion of ether lipids, achieved via knockout of the fatty acid reductase 1 (FAR1) enzyme^23^ (Supplementary Fig. 4b), also resulted in a significant decrease in intracellular iron levels, in this case in the OVCAR8 human high-grade serous ovarian cancer cell line, which exhibits mesenchymal characteristics^40^ (Fig. 2e, 2f). Furthermore, we noted that treatment of AGPS KO cells with ferric ammonium citrate (FAC)^41, 42^, which provides an exogenous source of ferric ions, re-sensitized cultured AGPS KO mesenchymal breast and ovarian carcinoma cells to ferroptosis induction (that could be rescued by the ferroptosis inhibitor liproxstatin-1), doing so even in the absence of elevated ether phospholipids (Fig. 2g, 2h).

We further extended this analysis by studying the behavior of mammary carcinoma tumors in vivo. Consistent with our in vitro data, ICP-MS indicated that total iron levels are reduced in breast tumors derived from implanted pB3 AGPS KO cells relative to those arising from either pB3 WT or pB3 AGPS addback cells (Fig. 2i). These observations reinforced the notion that ether lipids are critical regulators of intracellular iron levels, a rate-limiting component of ferroptosis^7^.

### Ether lipids facilitate CD44-mediated iron endocytosis

We proceeded to investigate the mechanism(s) by which membrane-associated ether lipids could regulate intracellular iron levels. This led us to examine the behavior of two proteins that are known to act as major mediators of cellular iron import— transferrin receptor 1 (TfR1)^43^ and CD44^35^—and whether their functioning was altered in response to loss of ether phospholipids. While CD44 is best known as a cell-surface cancer stem-cell marker^44, 45^, recent research has revealed its critical role in mediating endocytosis of iron-bound hyaluronates in CSC-enriched cancer cells and in activated immune cells^35, 39^. To monitor these two alternative iron import mechanisms, we performed endocytosis kinetics experiments using fluorescently labeled transferrin as a proxy for TfR1 internalization and fluorescently labeled hyaluronic acid (HA)—whose main plasma membrane receptor is CD44—as a marker for CD44 internalization^46, 47^.

We observed that the rate of TfR1 endocytosis was marginally affected by ether lipid deficiency in pB3 cells (Fig. 3a, Supplementary Fig. 5a). In stark contrast, CD44mediated endocytosis was significantly impaired in these ether lipid-deficient cancer cells (Fig. 3b, Supplementary Fig. 5b). Conversely, CD44-dependent iron import in AGPS KO cells could be restored to WT levels upon AGPS addback (Fig. 3b, Supplementary Fig. 5b). Reduction in the rate of internalization of CD44 but not TfR1 was also observed in ether lipid-deficient PyMT-1099 TGF-β-treated cells (Fig. 3c, 3d, Supplementary Fig. 5c, 5d). Therefore, ether lipids play a critical role in modulating intracellular iron levels by regulating endocytosis of CD44 but not transferrin receptor. These findings were consistent with our previous observations that TfR1 and CD44 localize to distinct endocytic vesicles in CSC-enriched cancer cells^35, 48^, making plausible that their internalization was governed by distinct endocytic mechanisms.

**Fig. 3:**
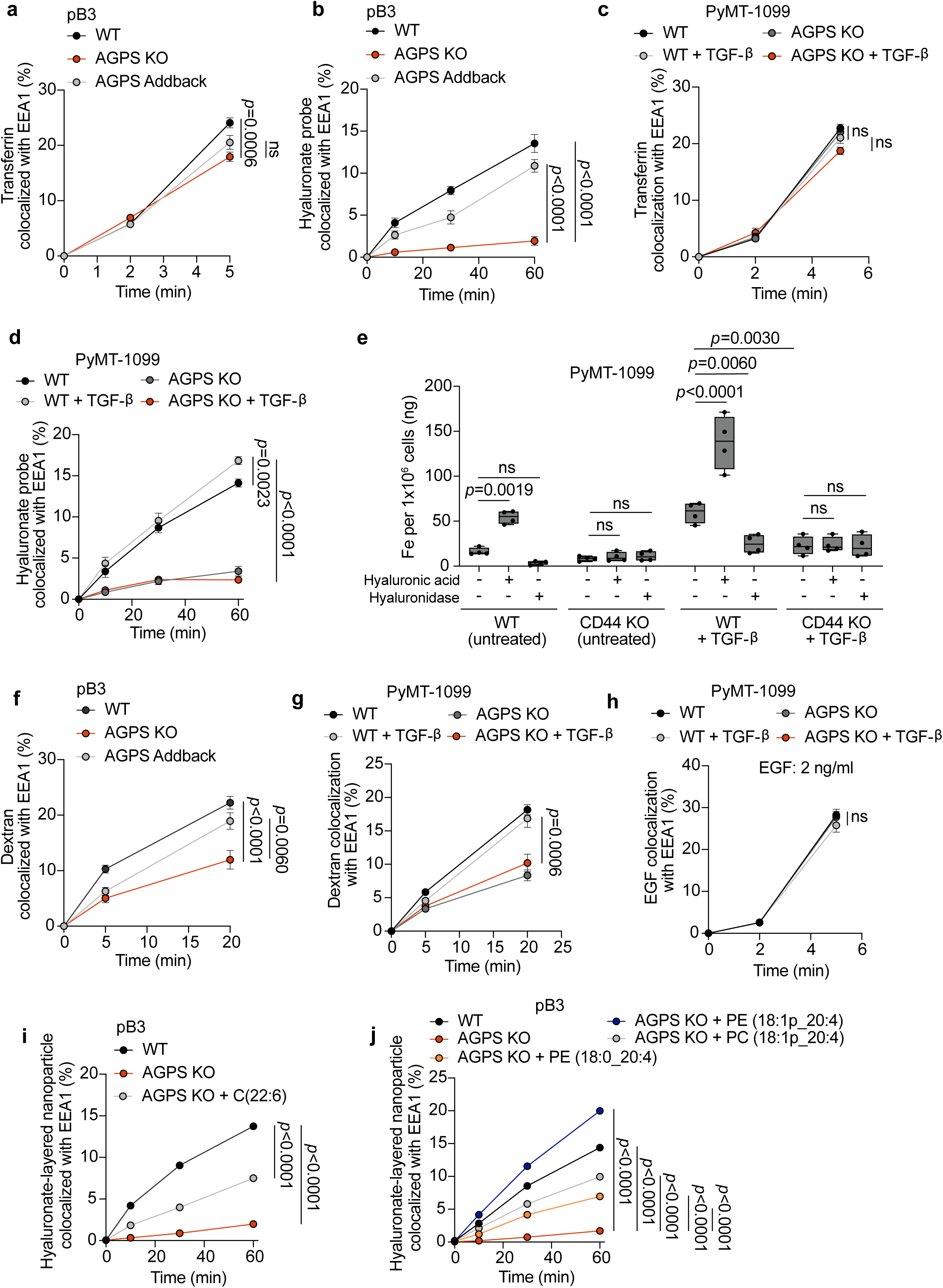
Ether lipids facilitate CD44-mediated iron endocytosis. a. Endocytic transport of fluorescently labeled transferrin as assessed by quantitative colocalization with an early endosomal marker (EEA1) in pB3 WT, AGPS KO, and AGPS addback cells. b. Endocytic transport of fluorescently labeled hyaluronate probe as assessed by quantitative colocalization with an early endosomal marker (EEA1) in pB3 WT, AGPS KO, and AGPS addback cells. c. Endocytic transport of fluorescently labeled transferrin as assessed by quantitative colocalization with an early endosomal marker (EEA1) in PyMT-1099 WT or AGPS KO cells +/− pretreatment with TGF-β. d. Endocytic transport of fluorescently labeled hyaluronate probe as assessed by quantitative colocalization with an early endosomal marker (EEA1) in PyMT-1099 WT or AGPS KO cells +/− pretreatment with TGF-β. e. Inductively coupled plasma-mass spectrometry (ICP-MS) of cellular iron following treatment with either hyaluronic acid or hyaluronidase in PyMT-1099 WT or CD44 KO cells +/− pretreatment with TGF-β (n=4). f. Endocytic transport of dextran as assessed by quantitative colocalization with the early endosomal marker EEA1 in pB3 WT, AGPS KO, and AGPS addback cells. g. Endocytic transport of dextran as assessed by quantitative colocalization with the early endosomal marker EEA1 in PyMT-1099 WT or AGPS KO cells +/− pretreatment with TGF-β. h. Endocytosis of EGFR as assessed by quantitative colocalization of internalized fluorescently labeled EGF with an early endosomal marker (EEA1) in PyMT-1099 WT or AGPS KO cells +/− pretreatment with TGF-β. Cells were treated with 2 ng/ml Alexa 555-conjugated EGF. i. Endocytic transport of hyaluronate-layered nanoparticle assessed by quantitative colocalization of internalized nanoparticles with an early endosomal marker (EEA1) in pB3 WT or AGPS KO cells +/− pretreatment (16-18 hr) with polyunsaturated fatty acid (PUFA) BSA conjugate C(22:6). j. Endocytic transport of hyaluronate-layered nanoparticle assessed by quantitative colocalization of internalized nanoparticles with an early endosomal marker (EEA1) in pB3 WT or AGPS KO +/− pretreatment (16-18 hr) with liposomes composed of the following: PE (18:0_20:4), PE (18:1p_20:4), and PC (18:1p_20:4). All data shown as mean +/− SEM and statistical significance was calculated by one-way ANOVA with Tukey’s multiple comparisons test; ns, not significant. For all endocytosis assays n=10 fields were examined for each timepoint, and data are representative of two independent experiments. In some cases, error bars are smaller than the symbol size and not visible. Panels a, b & f: pB3 WT and AGPS KO cells were transduced with the respective vector control plasmids.

To further support the role of CD44 in promoting iron uptake—acting via endocytosis of HA—we demonstrate that knocking out the gene encoding CD44 or, alternatively, treating cancer cells with the HA-degrading enzyme, hyaluronidase, led to a reduction in intracellular iron levels (Fig. 3e). Conversely, supplementing WT cells with HA significantly increased intracellular iron levels (Fig. 3e). Similar observations were seen in human OVCAR8 cells (Supplementary Fig. 7f). Taken together, these observations further supported the central role of CD44 in mediating iron uptake in these cancer cells^35^.

We then studied whether the defect in CD44 endocytosis observed in ether lipid-deficient cells was limited specifically to CD44 or other cell-surface proteins internalized by a similar mechanism. CD44 is known to undergo a type of clathrin- and dynamin-independent form of endocytosis^48–50^. To test whether loss of ether phospholipids had a wider effect on the clathrin- and dynamin-independent mode of endocytosis, we examined the rate of uptake of dextran (70 kDa), a branched polysaccharide known to undergo endocytosis by a clathrin-independent mechanism^49^ (pinocytosis). Similar to CD44, we observed that loss of AGPS also induced a significant reduction in the rate of dextran endocytosis; this behavior could be reversed by restoration of ether phospholipid levels achieved by AGPS addback (Fig. 3f, 3g, Supplementary Fig. 5e, 5f).

The internalization mechanism of CD44 and dextran differs from that of many plasma-membrane proteins, including TfR1 and EGFR, which undergo clathrin-dependent endocytosis, in which small invaginations of clathrin-coated pits undergo scission facilitated by the GTPase dynamin^49, 51^. As predicted by the distinct actions of these two mechanisms, we observed that loss of ether lipids had a negligible effect on the rate of EGFR endocytosis (Fig. 3h, Supplementary Fig. 6a, 7a-b). Taken together, these observations indicate that internalization of extracellular and cell-surface molecules is mediated by distinct mechanisms which differ in their dependence on plasma membrane ether phospholipids.

To further characterize the clathrin-independent mechanism used by CD44 and dextran, we next pursued both genetic and pharmacologic approaches that target against either the CLIC/GEEC pathway or pinocytosis. We found that inhibition of the CLIC/GEEC pathway via siRNA-mediated knockdown of PICK1^52^ or treatment with 7keto-cholesterol^53, 54^ significantly reduced CD44 internalization, which in turn led to decreased total cellular and lysosomal iron (Supplementary Fig. 8a-8c, 8e, 8g, 8h). We also show that blocking pinocytosis via knockdown of CtBP1^55–57^ or treatment with Cytochalasin-D^58^ is capable of impairing CD44 internalization (albeit, to a lesser extent than CLIC/GEEC inhibition), resulting in lower but non-significant reductions in total cellular and lysosomal iron (Supplementary Fig. 8a-8c, 8d, 8f, 8h). These findings suggest that CD44 internalization depends more on the CLIC/GEEC pathway than on pinocytosis for iron delivery into cells.

Given that polyunsaturated phospholipids have been shown to promote endocytosis^59^, we investigated whether treatment with exogenous polyunsaturated fatty acids (PUFAs) could compensate for loss of ether lipids in regulating CD44 endocytosis. We observed that supplementation with exogenous PUFAs could not fully rescue impaired CD44 endocytosis induced by loss of ether lipids (Fig. 3i, Supplementary Fig. 6b). Furthermore, while liposomes containing ester-linked PUFA-PE phospholipids could only partially rescue CD44 endocytosis, ether-linked PUFA-PE or -PC liposomes achieved an even greater degree of rescue, with ether-linked PUFA-PE liposomes fully restoring CD44 endocytosis to levels exceeding those of WT cells (Fig. 3j, Supplementary Fig. 6c). These findings demonstrate that both the presence of the ether linkage in phospholipids as well as the type of headgroup present within phospholipids play important roles in supporting efficient CD44-mediated endocytosis.

### Ether lipid deficiency impairs membrane biophysical properties

The above observations did not provide mechanistic insights into how changes in the composition of membrane ether lipids could exert an effect on CD44 internalization. As observed by others, non-clathrin-mediated endocytosis, which is employed by CD44, is particularly sensitive to changes in the physicochemical properties of the lipid bilayer forming plasma membranes^50, 60–65^. Such changes can influence membrane tension and fluidity and the stability and formation of lipid rafts, all of which impact the assembly and dynamics of clathrin-independent, cell-surface endocytic structures. Hence, we hypothesized that ether lipids alter the biophysical properties of the plasma membrane to facilitate elevated iron endocytosis via CD44. To substantiate these hypotheses, we quantified the abundance of ether lipids in the plasma membrane. Lipidomics profiling of pB3-derived giant plasma membrane vesicles (GPMV) showed that ether lipids make up ∼23% of plasma membrane lipids and are depleted to ∼5% upon AGPS KO (Supplementary Fig. 9a-f).

Alterations in membrane tension have long been demonstrated to affect endocytosis^66–71^. This prompted us to assess the effects of ether lipid deficiency on plasma membrane tension. Membrane tension measures the forces exerted on a defined cross-section of the plasma membrane, which is influenced by both the in-plane tension of the lipid bilayer and the attachment of the plasma membrane to the underlying cell cortex^72, 73^.

To quantify membrane tension directly, we generated a membrane tether using an optically trapped bead and measured the pulling force ( ) and the tube radius (R) to calculate membrane tension (σ) of living cells^59, 74^ (Fig. 4a, 4b). We found that depletion of ether phospholipids led to a significant increase in membrane tension in pB3 AGPS KO cells relative to the corresponding pB3 WT cells (Fig. 4c). This shift was largely attenuated upon restoration of AGPS expression in pB3 AGPS KO cells or upon exposure of cultured cells to liposomes composed of ether phospholipids (Fig. 4c). Consistent with earlier measurements, treatment of pB3 AGPS KO cells with ether PUFA-PE liposomes increased the rate of CD44 endocytosis to levels comparable to those of pB3 WT cells. In contrast, treatment of pB3 AGPS KO cells with liposomes derived from monounsaturated phosphatidylethanolamine (MUFA-PE) ether lipids only partially rescued CD44 endocytosis (Fig. 4d, Supplementary Fig. 6d). No changes were observed in the rate of clathrin-dependent TfR1 endocytosis under these conditions (Fig. 4e, Supplementary Fig. 6e) and loss of CD44 showed modest effects on membrane tension (Supplementary Fig. 7d-7e). Taken together, these results provide evidence that ether lipids facilitate CD44-mediated iron endocytosis in cancer cells, in part by decreasing membrane tension.

**Fig. 4:**
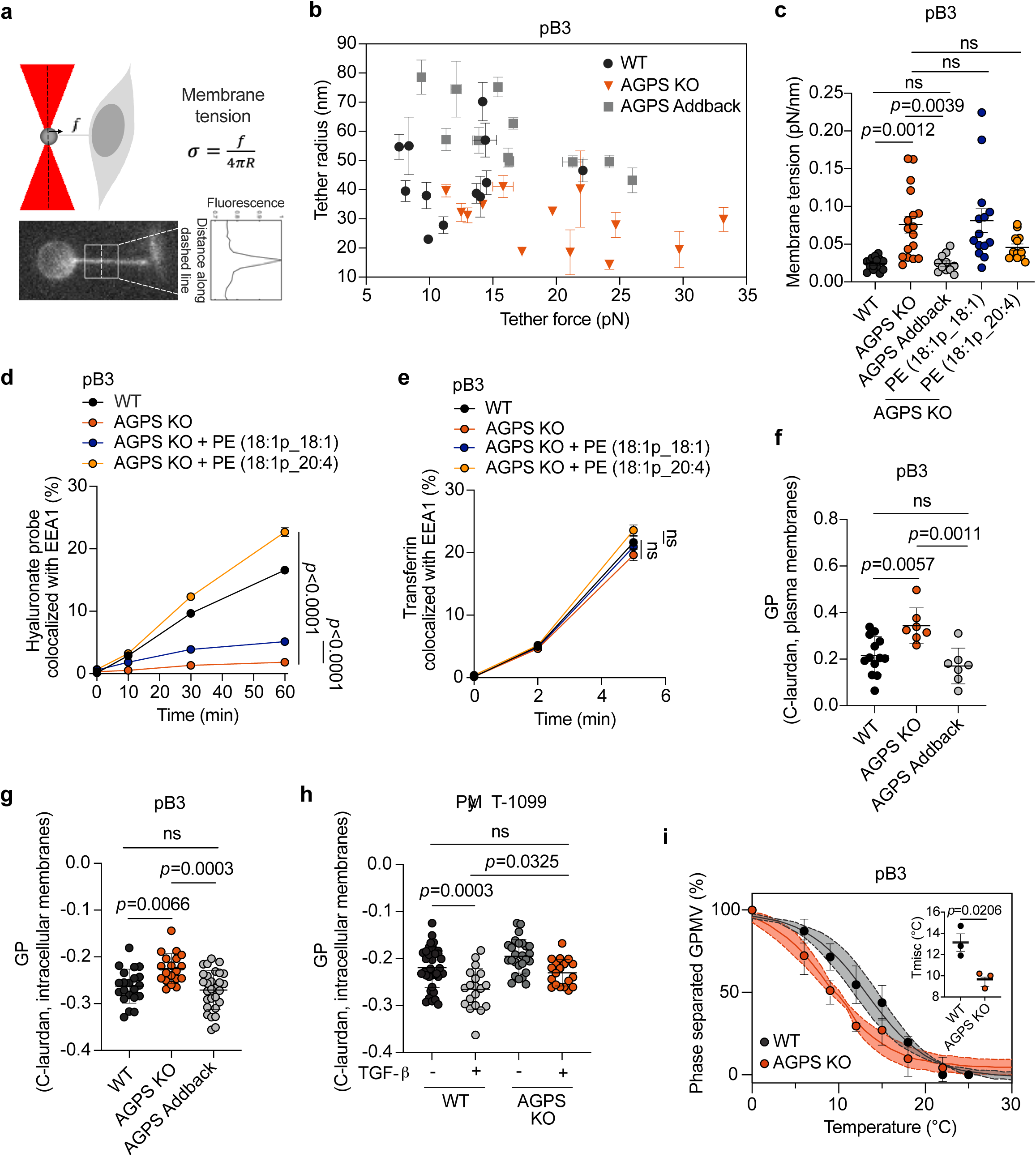
Ether lipid deficiency impairs membrane biophysical properties. a. Schematic of membrane tether pulling assay and fluorescence image showing a tether pulled from the plasma membrane of a pB3 cell using an optically trapped 4 µm anti-Digoxigenin coated polystyrene bead. b. Graph showing tether radius (R) and tether force measurements ( ) in pB3 WT, AGPS KO, and AGPS addback cells. All data shown as mean +/− SD. c. Membrane tension measurements in pB3 WT, AGPS KO pretreated with the indicated ether phospholipid liposomes, and AGPS addback cells. All data shown as mean +/− SEM. d. Endocytic transport of fluorescently labeled hyaluronate probe as assessed by quantitative colocalization with an early endosomal marker (EEA1) in pB3 WT or AGPS KO cells pretreated with the indicated ether phospholipid liposomes. All data shown as mean +/− SEM. e. Endocytic transport of fluorescently labeled transferrin as assessed by quantitative colocalization with an early endosomal marker (EEA1) in pB3 WT or AGPS KO cells pretreated with 20µM of the indicated ether phospholipid liposomes. All data shown as mean +/− SEM. f. Generalized polarization (GP) values of C-laurdan-labeled plasma membranes from pB3 WT, AGPS KO and AGPS addback cells. Data is shown as mean GP +/− SD. g. GP values of C-laurdan-labeled intracellular membranes from pB3 WT, AGPS KO and AGPS addback cells. Data is shown as mean GP +/− SD. h. GP values of C-laurdan-labeled intracellular membranes from PyMT-1099 WT or AGPS KO cells treated with or without 2 ng/ml TGF-β for 10 d. Data shown as mean GP +/− SD. i. Representative giant plasma membrane vesicle (GPMV) phase separation curves from AGPS KO versus WT pB3 cells. Curves were generated by counting ≥ 20 vesicles/temperature/condition at >4 temperatures. The data was fit to a sigmoidal curve to determine the T_misc_. Data shown as the mean +/− SD of 3 independent experiments with the average fit shown. Inset shows mean T_misc_ +/− SD upon loss of AGPS in pB3 cells. Statistical significance was calculated using unpaired, two-tailed t-test. Unless otherwise noted statistical significance was calculated by one-way ANOVA with Tukey’s multiple comparisons test; ns, not significant. Examined n=10 fields for all endocytosis-related experiments and at two independent replicates were performed. For panels b, c, f & g: pB3 WT and AGPS KO cells were transduced with the respective vector control plasmids.

Membrane lipid packing can also impact endocytosis^75–77^. This is related to the fluidity or viscosity of the lipid bilayer, with higher lipid packing correlating with higher viscosity. This affects the ease with which proteins and lipids undergo lateral diffusion and conformational changes within a lipid bilayer, thereby affecting endocytosis-related signaling^59^. This prompted us to investigate the contribution of ether lipids to membrane lipid packing. We used C-laurdan, a lipid-based, polarity-sensitive dye, which yields a spectral emission shift dependent on the degree of lipid packing^78^. These measurements are used to calculate a unitless index, termed generalized polarization (GP), where a higher GP indicates increased lipid packing^78^. Our measurements using C-laurdan indicated that a reduction in ether lipid levels resulted in a measurable and significant increase in membrane packing (Fig. 4f-4h, Supplementary Fig. 7c) which, like increases in membrane tension, negatively affects membrane deformability^75^.

The association of CD44 with lipid rafts, which are dynamic plasma membrane nanodomains, is known to be critical for CD44-mediated HA endocytosis^79^. Previous work has shown that PUFA-containing lipids can stabilize lipid rafts by increasing the packing contrast between ordered and disordered phases^80–83^. Thus, we hypothesized that lipid raft stability might be reduced in cells lacking AGPS and therefore PUFA-containing ether lipids which would result in impairment of CD44 endocytosis^84^. Accordingly, we measured the miscibility transition temperature (T_misc_) which is hypothesized to be related to the stability membrane domains of isolated cellular plasma membranes. We observed a decrease in T_misc_ upon loss of AGPS in pB3 cells, which indicate a decrease in lipid raft stability (Fig. 4i). This finding supports a role for ether lipids in maintaining the plasma membrane organization through lipid raft microdomains, revealing yet another biophysical property of lipid bilayers that can influence CD44 endocytosis.

It is noteworthy that clathrin-independent endocytosis exhibits a greater dependency on the membrane biophysical properties assessed above^50, 60–65^. This may explain why loss of ether lipids can exert a significant effect on the rate of clathrin-independent, CD44-mediated iron endocytosis but negligible effects on the clathrin-dependent TfR1 endocytosis. Furthermore, these findings revealed a mechanism by which membrane-associated ether lipids could govern a major mechanism of iron internalization, which may impact the vulnerability of cancer cells to ferroptosis inducers.

### Loss of ether lipids decreases metastasis and cancer cell stemness

Prior studies have demonstrated that reduced membrane tension and elevated intracellular iron can promote cancer metastasis^85–89^. These findings prompted us to investigate whether changes in the ether lipid composition of cancer cells impact key steps of the multi-step invasion-metastasis cascade, notably extravasation efficiency, post-extravasation proliferation^90^ and importantly stemness as manifested by tumor-initiating ability.

We measured extravasation efficiency by employing an in vitro three-dimensional microvascular network system composed of human umbilical vein endothelial cells (HUVECs) and normal human lung fibroblasts. This system has been shown to accurately model some of the complex biological processes associated with cancer cell extravasation^91–95^. Using this defined experimental system, we found that loss of ether lipids significantly decreased extravasation efficiency (Fig. 5a-5c). Furthermore, we observed a strong reduction in overall metastatic burden following intracardiac injection in syngeneic hosts of the pB3 AGPS KO cells relative to corresponding WT cells (Fig. 5d-5f). We note that ether lipid deficiency had a modest effect on primary tumor growth kinetics (Fig. 5g-5i). A decrease in metastatic burden was also observed upon knockout or *AGPS* and *FAR1* in OVCAR8 cells, and upon loss of CD44 in pB3 cells (Supplementary Fig. 10a-10c).

**Fig. 5:**
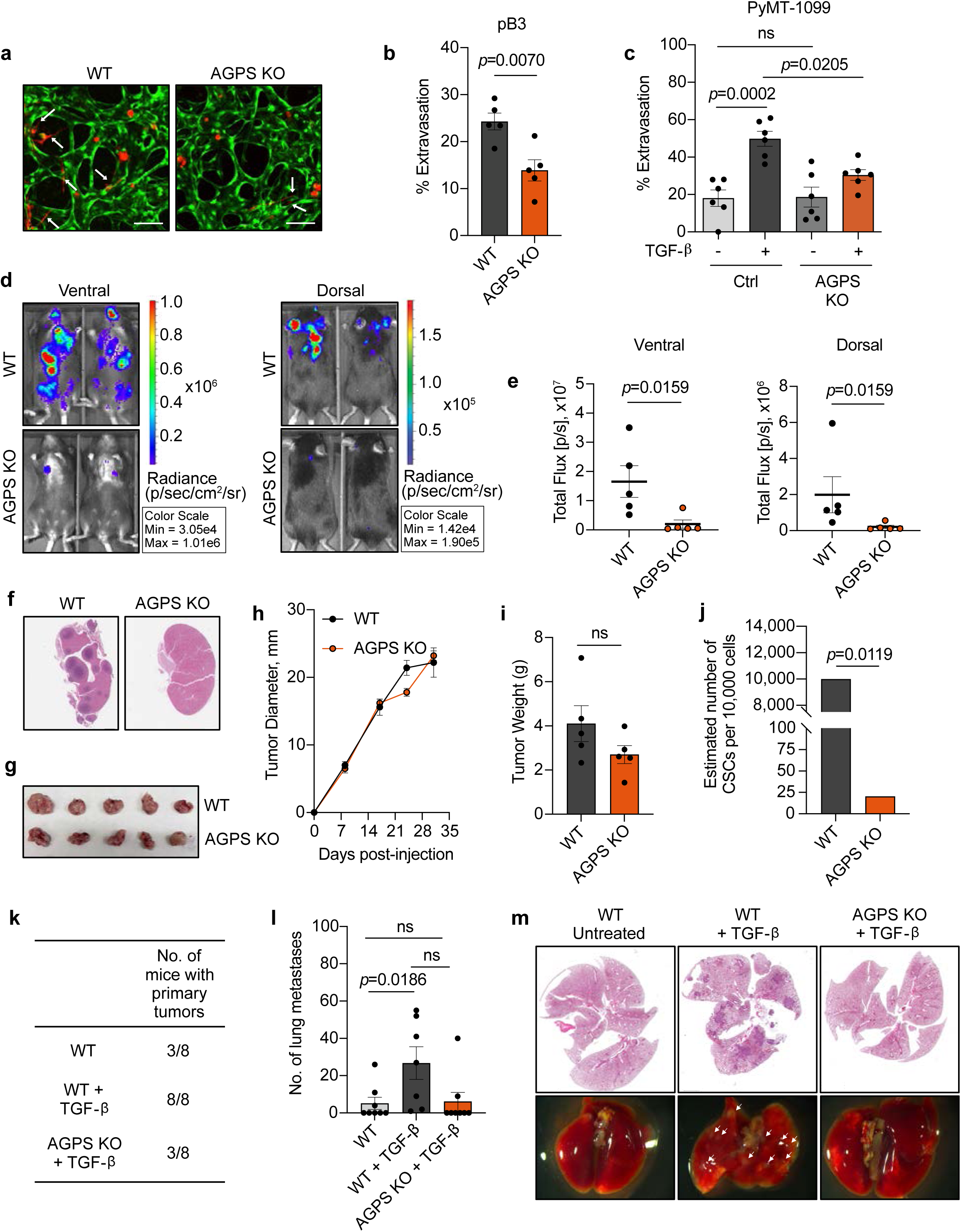
Loss of ether lipids decreases metastasis and cancer cell stemness. a. Representative confocal images of extravasated tdTomato-labeled pB3 WT and AGPS KO cells from an in vitro microvascular network (see methods), over 24 h. Data representative of two independent replicates (scale bar = 100 µm). b. Quantification of extravasated pB3 WT and AGPS KO cells from microvascular network. Data represent the mean percentage of extravasated cells per device +/− SEM (n=5). Significance was calculated using unpaired, two-tailed t-test. Data representative of two independent replicates. c. Quantification of extravasated tdTomato-labeled PyMT-1099 cell line derivatives from microvascular network. Data represent the mean percentage of extravasated cells per device +/− SEM (n=6). Significance was calculated by one-way ANOVA with Tukey’s multiple comparisons test; ns, not significant. Data representative of two independent replicates. d. Representative in vivo imaging system (IVIS) images of overall metastatic burden in C57BL/6 female mice following intracardiac injection of pB3 WT (n=5) and AGPS KO (n=5) cells. e. Quantification of metastatic burden in C57BL/6 female mice following intracardiac injection of pB3 WT and AGPS KO cells (n=5). Graph shows the mean +/− SEM and statistical significance was calculated using two-tailed Mann-Whitney test. f. Representative images of H&E-stained sections of harvested kidneys from C57BL/6 female mice following intracardiac injection of pB3 WT or pB3 AGPS KO cells. g. Gross images of primary tumors derived from pB3 WT control cells and pB3 AGPS KO cells. h. Tumor growth kinetics of primary tumors derived from pB3 WT control cells and pB3 AGPS KO cells (n=5 mice per group). Data shown as mean +/− SEM. i. Bar graph showing the average weight of primary tumors derived from pB3 WT and AGPS KO cells (n=5). Graph shows the mean +/− SEM and statistical significance was calculated using two-tailed Mann-Whitney test; ns, not significant. j. Estimated number of cancer stem cells (CSCs) per 10,000 cells as calculated by extreme limiting dilution analysis (ELDA) software. Tumor-initiating capacity was assessed following implantation of pB3 WT or pB3 AGPS KO cells into the mammary fat pad of C57BL/6 mice. *P* values, two-sided χ2 pairwise test. k. Table showing the number of mice with palpable primary tumors at 121 d post orthotopic implantation of PyMT-1099 WT or AGPS KO cells +/− pretreatment with TGF-β into female NSG mice. l. Quantification of lung metastases for aforementioned experiment. Data represents mean number of lung metastases +/− SEM (n=8 for WT and AGPS KO + TGF-β, n=7 for WT + TGF-β). Statistical significance was calculated by one-way ANOVA with Tukey’s multiple comparisons test; ns, not significant. m. Representative images of H&E-stained lungs harvested from female NSG mice from aforementioned experiment. Arrows indicate metastases.

Given that high CD44 expression and elevated intracellular iron levels are positively correlated with cancer cell stemness^35, 44, 96^, we investigated whether ether lipid deficiency could modulate CSC properties, by employing mammosphere formation assays^97^ and experimental limiting dilution tumor-implantation studies. These experiments indicated that loss of ether lipids in pB3 cells decreases cancer cell stemness (Fig. 5j, Supplementary Fig. 10d-10e), which, as we have found in other investigations, serve as a reliable marker of metastasis-initiating capacity^98^.

Additionally, we show that treatment of human MCF7 breast adenocarcinoma cells with oncostatin M (OSM) increases the sub-population CD44^hi^/CD24^lo^ cells (Supplementary Fig. 11a, 11b), resulting in elevated PUFA-containing ether phospholipids and increased intracellular iron (Supplementary Fig. 11c-11g). This is consistent with our TGFβ-treated PyMT-1099 model (Supplementary Fig. 12a-12c).

Moreover, we found that loss of ether lipids significantly attenuates the tumor-initiating potential and metastatic capacity of PyMT-1099 AGPS KO TGF-β-treated cells following orthotopic implantation into the mammary stromal fat pad compared to PyMT1099 WT TGF-β-treated cells (Fig. 5k-5m). Collectively, our findings indicate that ether lipids play critical roles in promoting cancer cell stemness and resulting post-extravasation colonization.

## DISCUSSION

Our findings underscore the importance of integrating membrane biophysical properties into existing genetic and biochemical frameworks for understanding cancer cell fate. While phospholipids have long been implicated in cellular transformation, the focus thus far has primarily been to study the actions of specific lipids (e.g. inositol phospholipids) and their signaling derivatives^99^. Our study reveals a distinct and underappreciated mechanism by which lipids influence malignancy. Specifically, we identify ether lipids as key modulators of plasma membrane biophysical properties that regulate iron uptake and neoplasia-related phenotypes such as metastasis and cancer cell stemness/tumor-initiating potential. Importantly, this biophysical configuration creates a unique vulnerability of cancer cells to ferroptosis and suggests that targeting lipid metabolism and iron homeostasis could be exploited to suppress subpopulations of highly metastatic and drug-tolerant carcinoma cells^7, 100^.

Ether phospholipids have been widely characterized as participants in ferroptosis through their role as substrates prone to iron-catalyzed oxidation^22^. However, our findings indicate an entirely distinct biochemical mechanism is operative here, whereby ether lipids directly modulate the levels of intracellular iron, a rate-limiting parameter governing ferroptosis susceptibility^4, 35, 86^. By emphasizing the role of membrane biophysical properties in mediating iron uptake, we depart from the conventional portrayal of phospholipids simply as substrates of oxidation. This shift in perspective has the potential to open new avenues for research, as it challenges researchers to explore the biophysical aspects of membranes as a new dimension in the regulation of this cell death program.

Although most lipid classes remained largely unchanged upon AGPS KO, it is plausible that subtle changes in membrane lipid composition beyond ether lipid levels may also be responsible for the phenotypes described here. Moreover, in addition to perturbed membrane receptor dynamics promoted by changes in membrane tension, the observed decrease in metastatic capacity of AGPS KO cells may also result from changes in oncogenic signaling lipids^24^, mechanisms that are not addressed here.

Alterations in intracellular iron level can impact gene expression via various mechanisms, including modulation of chromatin-modifying enzyme activity^35, 101, 102^. For example, an increase of intracellular iron levels has been shown to promote the activity of iron-dependent histone demethylases^35, 101, 102^, impacting gene expression profiles underlying cell plasticity^35^ and immune cell activation^39^. Our finding that ether lipid deficiency reduces intracellular iron levels explains, at least in part, how loss of ether lipids may affect cancer-associated transcriptional programs, acting at the epigenetic level and enabling a variety of malignancy-associated cell phenotypes including metastasis and cancer stemness. Such mechanisms may act in concert with processes independent from iron, which are also regulated by ether lipids to affect cancer malignancy traits.

In the longer term, the implications of our findings may reach well beyond cancer pathogenesis. Thus, we suggest that the interplay between membrane biophysics and iron biology could represent a fundamental determinant of cell fate, influencing diverse processes such as differentiation, immune activation, wound healing, and embryonic development.

## METHODS

### Cell lines

The pB2 and pB3 MMTV-PyMT-derived murine breast cancer cell lines were a kind gifts from the laboratory of Harold L. Moses^32^. 687g cells (also called EpCAM^Lo^Snail-YFP^Hi^) were originally established from tumors that developed in the MMTV-PyMT-Snail-IRESYFP reporter mouse model, previously developed by the Weinberg lab^103^. pB3 and 687g cell lines were cultured in 1:1 DMEM/F12 medium containing 5% adult bovine serum with 1% penicillin-streptomycin and 1% non-essential amino acids^32^. The PyMT-1099 murine breast cancer cell line was a kind gift from the laboratory of Gerhard Christofori and cultured in DMEM supplemented with 10% fetal bovine serum, 1% penicillinstreptomycin and 1% glutamine^29^. For select experiments, these cells were treated with 2 ng/ml of TGF-β for 10 days prior to performing subsequent analyses. OVCAR8 cells were obtained from the laboratory of Joan Brugge and cultured in 1:1 MCDB 105 medium/Medium 199 Earle’s Eagles medium supplemented with 10% fetal bovine serum and 1% penicillin-streptomycin. MCF7 (ATCC, HTB-22) cells were cultured in Dulbecco’s Modified Eagle Medium GlutaMAX (DMEM, ThermoFisher Scientific, 61965059) supplemented with 10% Fetal Bovine Serum and Penicillin-Streptomycin mixture (BioWhittaker/Lonza, DE17-602E) and treated with oncostatin M (OSM, R&D systems, 295-OM-050, 100 ng/mL, 72 h). All cells were cultured in a humidified incubator at 37°C with 5% CO_2_. All cells were negative for mycoplasma. Human cell line authentication (CLA) analysis of OVCAR8 cells were performed by the Duke University DNA Analysis facility. Established murine lines have not been STR profiled.

### Animal studies

All animal experiments were performed in the animal facility at the Koch Institute for Integrative Cancer Research at MIT. Mice we housed under a 12-h light/12-h dark cycle, temperatures in the range 20–22 °C and humidity between 30–70%. Animal experiments performed in this study received approval by the MIT Institutional Animal Care and Use Committee. Tumor burden, where measurable, were not allowed to exceed 1 cm^2^ unless approved by the Institutional Animal Care and Use Committee. Animals showing discomfort or distress were humanely euthanized following the MIT Institutional Animal Care and Use Committee guidelines. For primary tumor growth studies: 1 million cells were resuspended in 20% Matrigel/PBS and injected into the mammary fat pad of 6-8 week old female mice. C57BL/6 mice (Jackson Laboratories; strain name: C57BL/6J; stock number: 000664) were used for in vivo experiments with pB3 cells. NSG mice (Jackson Laboratories; strain name: NOD.Cg-Prkdc^scid^ Il2rg^tm1Wjl^/SzJ; stock number: 005557) were used for in vivo experiments with PyMT1099 cells. These cells were pretreated with TGF-β (2 ng/ml) for 10 days prior to injection. Tumor size was measured once a week using a vernier caliper and tumor volume was calculated using the formula: Tumor volume = (length x width^2^)/2, where length represents the largest tumor diameter, and width represents the perpendicular tumor diameter. For limiting dilution tumor-initiating assays, pB3 WT or pB3 AGPS KO cells were resuspended in 20% Matrigel/PBS and injected into the mammary fat pad of 6-8 weeks old female C57BL/6 mice at the following dilutions: 100,000, 10,000, 1000, 100 cells. Animals were assessed for palpable tumors after 39 days post injection. The estimated number of CSCs was calculated using the extreme limiting dilution analysis (ELDA) software^104^. For experimental metastasis involving pB3 cell lines, 0.2 million GFP-luciferized cells were resuspended in 200µl of PBS and injected into the left ventricle of 6-8 weeks old female C57BL/6 mice. Metastatic burden was measured after 10 d post-injection via bioluminescence in live animals using the spectrum in vivo imaging system (IVIS)^105^. Images were analyzed using Living Image software (PerkinElmer, version 4.8.2). For OVCAR8 cells, 1.5 million cells were resuspended in PBS and implanted into 6-8 weeks old female athymic nude mice (Jackson Laboratories; strain name: NU/J; stock number: 002019) via intraperitoneal injections. Metastatic burden was assessed after 6 weeks using a fluorescence dissecting microscope.

### Generation of gene-edited cell lines using CRISPR/Cas9

Except for OVCAR8 cells, all AGPS KO single-cell clones were generated via transient transfection with mouse AGPS CRISPR/Cas9 KO Plasmids (Catalog no. sc-432759, Santa Cruz) according to the manufacturer’s instructions. GFP-positive cells were sorted via fluorescence-activated cell sorting (FACS) into 96-well plates with one cell per well and single-cell clones were subsequently expanded. AGPS KO single-cell clones were assessed for loss of AGPS expression via western blot analysis. pB3 and PyMT-1099 AGPS addback cells were generated by transducing an AGPS KO singlecell clone with pLV[Exp]-Puro-EF1A-mAgps lentiviral vector (VectorBuilder). pB3 and PyMT-1099 AGPS KO cells expressing pLV[Exp]-Puro-EF1A-Stuffer_300bp (VectorBuilder) were established as controls for AGPS addback experiments and noted in the figure legends where used. Lentivirus was produced by transfecting HEK293T cells with viral envelope (VSVG, Addgene) and packaging plasmids (psPAX2, Addgene). Viral supernatant was collected after 48 h and filtered through a 0.45µm filter. Stably transduced cells were selected with 2 µg/ml puromycin. OVCAR8 FAR1 KO and AGPS KO single cell clones as well as nontargeting control cells were established as previously described^22^. pB3 CD44 KO cells (bulk) were generated using human CD44 CRISPR/Cas9 KO Plasmids (Catalog no. sc-419558, Santa Cruz) according to the manufacturer’s instructions. After 48 h post-transfection, cells were sorted by flow cytometry for GFP positive cells, expanded in culture, and re-sorted twice for CD44 negative cells using Alexa Fluor® 647 anti-mouse/human CD44 Antibody. Cells were maintained as bulk CD44 KO cells.

**Supplementary Table 1:** Plasmid information. The table below indicates the plasmids used to generate gene-edited cell lines.

| Plasmid | Source |
| --- | --- |
| pLV[Exp]-Puro-EF1A-Agps (mouse) | VectorBuilder |
| pLV[Exp]-Puro-EF1A-Stuffer_300bp | VectorBuilder |
| Mouse AGPS CRISPR/Cas9 KO Plasmids | Catalog no. sc-432759, Santa Cruz |
| Mouse CD44/HCAM CRISPR/Cas9 KO Plasmids | Catalog no. sc-419558, Santa Cruz |
| LentiCRISPRv2-puro-Nontargeting | Published<sup>22</sup> |
| LentiCRISPRv2-puro-human AGPS sg | Published<sup>22</sup> |
| LentiCRISPRv2-puro-human FAR1 sg | Published<sup>22</sup> |
| pLV-EF1A-eGFP-LUC | VectorBuilder |
| pCDH-EF1-Luc2-P2A-tdTomato | Plasmid #72486, Addgene |

### Lipidomics analysis

#### Sample preparation (whole cell preparations)

Briefly, 1×10^6^ cells were plated in individual wells of a 6 well dish. No more than 16 hours later, cells were washed with DPBS (Dulbecco’s PBS without Mg and Ca), trypsinized, and washed twice again with before resuspending in 300uL of DPBS and storing at -80°C until further processing.

#### Sample preparation (GPMV preparations)

Giant plasma membrane vesicles (GPMVs) were isolated from pB3 WT, AGPS KO, and AGPS addback, as previously described with minor modifications^106^. Briefly, three 10cm dishes of approximately 70% confluency prepared for each cell line were washed once with phosphate-buffered saline (PBS) and twice with GPMV buffer (10mM HEPES, 150mM NaCl, 2mm CaCl2 – pH 7.4). Fresh N-ethylmaleimide (Millipore Cat. No. E3876-5G) was added to GPMV buffer to yield a final concentration of 5 mM, then 4mL of this buffer was added to each culture dish followed by incubation for 3 hours at 37°C. Supernatants containing GPMVs were then collected and pooled for each condition and passed through 5 µm centrifugal filters (Millipore Sigma Cat. No. UFC30SV00) by centrifuging at 500 xg to remove any whole cells. The pooled filtrates were subsequently centrifuged at 20,000 × g for 2 hours at 4°C and resuspended in 300 µL of DPBS. The presence of vesicles was confirmed by light microscopy before storing at 80°C until further processing. Three independent biological replicates were collected and used for lipidomic analysis.

#### Lipid extraction for mass spectrometry lipidomics

Mass spectrometry-based lipid analysis was performed by Lipotype GmbH (Dresden, Germany) as described^107^. Lipids from whole cells or isolated GPMVs were extracted using a chloroform/methanol procedure^108^. Samples were spiked with internal lipid standard mixture containing: cardiolipin 14:0/14:0/14:0/14:0 (CL), ceramide 18:1;2/17:0 (Cer), diacylglycerol 17:0/17:0 (DAG), hexosylceramide 18:1;2/12:0 (HexCer), lysophosphatidate 17:0 (LPA), lyso-phosphatidylcholine 12:0 (LPC), lysophosphatidylethanolamine 17:1 (LPE), lyso-phosphatidylglycerol 17:1 (LPG), lysophosphatidylinositol 17:1 (LPI), lyso-phosphatidylserine 17:1 (LPS), phosphatidate 17:0/17:0 (PA), phosphatidylcholine 17:0/17:0 (PC), phosphatidylethanolamine 17:0/17:0 (PE), phosphatidylglycerol 17:0/17:0 (PG), phosphatidylinositol 16:0/16:0 (PI), phosphatidylserine 17:0/17:0 (PS), cholesterol ester 20:0 (CE), sphingomyelin 18:1;2/12:0;0 (SM), triacylglycerol 17:0/17:0/17:0 (TAG). After extraction, the organic phase was transferred to an infusion plate and dried in a speed vacuum concentrator. The dry extract was re-suspended in 7.5 mM ammonium formate in chloroform/methanol/propanol (1:2:4, V:V:V). All liquid handling steps were performed using Hamilton Robotics STARlet robotic platform with the Anti Droplet Control feature for organic solvents pipetting.

#### MS data acquisition

Samples were analyzed by direct infusion on a QExactive mass spectrometer (Thermo Scientific) equipped with a TriVersa NanoMate ion source (Advion Biosciences).

Samples were analyzed in both positive and negative ion modes with a resolution of R_m/z=200_=280000 for MS and R_m/z=200_=17500 for MSMS experiments, in a single acquisition. MSMS was triggered by an inclusion list encompassing corresponding MS mass ranges scanned in 1 Da increments^109^. Both MS and MSMS data were combined to monitor CE, DAG and TAG ions as ammonium adducts; LPC, LPC O-, PC, PC O-, as formate adducts; and CL, LPS, PA, PE, PE O-, PG, PI and PS as deprotonated anions. MS only was used to monitor LPA, LPE, LPE O-, LPG and LPI as deprotonated anions; Cer, HexCer and SM as formate adducts.

#### Data analysis and post-processing

Data were analyzed with in-house developed lipid identification software based on LipidXplorer^110, 111^. Only lipid identifications with a signal-to-noise ratio >5, and a signal intensity 5-fold higher than in corresponding blank samples were considered for further data analysis. Cholesterol was removed from each data set before analysis. Simple imputation was performed by replacing missing values with 0.2 * the minimum non-zero value for each lipid assayed, a modification from previously described methods^112^. Relative mol% levels of individual lipid species were determined by dividing each pmol value by the sum pmol of all lipids detected in that sample and multiplying by 100. Total ether lipid (and other lipid classes presented) were calculated by summation of mol% values of each lipid within that class. Lipid fold changes were calculated by dividing the average mol% of each lipid species between groups and applying a Log2 transformation. Corresponding p-values were calculated using the students t-test.

### Oxidized lipidomics

#### Sample preparation

100,000 cells per condition were plated in 6-well plates 24 h prior to the experiment. For PyMT-1099, cells were treated with TGF-β (2 ng/mL) for 10 d. pB3 cells were treated with 500 nM RSL3, OVCAR8 cells with 2 µM ML210 and TGF-β-treated PyMT-1099 cells with 10 µM ML210 for 24 h. Cells were subsequently washed with PBS and then with 150 mM ammonium bicarbonate. Cells were then scraped and resuspended in 150 mM ammonium bicarbonate and centrifuged at 300 xg for 5 min. The supernatant was removed, and cells were resuspended in 1 mL of 150 mM ammonium bicarbonate. The solutions were centrifuged at 13,500 xg for 10 min and the supernatant was removed. 200 µL of 150 mM sodium bicarbonate was added to the pellet and samples were flash frozen in liquid nitrogen. Cells were prepared in 5 independent biological replicates and lipidomics analysis was performed on the same day for all the replicates.

#### Lipid extraction for mass spectrometry lipidomics

For lipidomics analysis, the 200 µL cell lysates were spiked with 1.4 μL of internal standard lipid mixture containing 300 pmol of phosphatidylcholine 17:0-17:0, 50 pmol of phosphatidylethanolamine 17:0-17:0, 30 pmol of phosphatidylinositol 16:0-16:0, 50 pmol of phosphatidylserine 17:0-17:0, 30 pmol of phosphatidylglycerol 17:0-17:0 and 30 pmol of phosphatidic acid 17:0-17:0 and subjected to lipid extraction at 4 °C, as previously described^113^. The sample was then extracted with 1 mL of chloroform-methanol (10:1) for 2 h. The lower organic phase was collected, and the aqueous phase was reextracted with 1 mL of chloroform-methanol (2:1) for 1 h. The lower organic phase was collected and evaporated in a SpeedVac vacuum concentrator. Lipid extracts were dissolved in 100 μL of infusion mixture consisting of 7.5 mM ammonium acetate dissolved in propanol:chloroform:methanol [4:1:2 (vol/vol)].

#### MS data acquisition

Samples were analyzed by direct infusion in a QExactive mass spectrometer (Thermo Fisher Scientific) equipped with a TriVersa NanoMate ion source (Advion Biosciences). 5 µL of sample were infused with gas pressure and voltage set to 1.25 psi and 0.95 kV, respectively. PC, PE, PEO, PCOx and PEOx were detected in the 10:1 extract, by positive ion mode FTMS as protonated adducts by scanning m/z= 580–1000 Da, at R_m/z=200_=280 000 with lock mass activated at a common background (m/z=680.48022) for 30 s. Every scan is the average of 2 micro-scans, automatic gain control (AGC) was set to 1E6 and maximum ion injection time (IT) was set to 50 ms. PG and PGOx were detected as deprotonated adducts in the 10:1 extract, by negative ion mode FTMS by scanning m/z= 420–1050 Da, at R_m/z=200_=280 000 with lock mass activated at a common background (m/z=529.46262) for 30 s. Every scan is the average of 2 microscans. Automatic gain control (AGC) was set to 1E6 and maximum ion injection time (IT) was set to 50ms. PA, PAOx, PI, PIOx, PS and PSOx were detected in the 2:1 extract, by negative ion mode FTMS as deprotonated ions by scanning m/z= 400–1100 Da, at R_m/z=200_=280 000 with lock mass activated at a common background (m/z=529.46262) for 30 s. Every scan is the average of 2 micro-scans, automatic gain control (AGC) was set to 1E6 and maximum ion IT was set to 50 ms. All data was acquired in centroid mode. All lipidomics data were analyzed with the lipid identification software, LipidXplorer^111^. Tolerance for MS and identification was set to 2 ppm. Data were normalized to internal standards.

### Immunoblotting

Cells were washed with ice-cold PBS and lysed in cell lysis buffer (Cell Signaling Technology, Cat. #9803S) containing 1mM PMSF protease inhibitor (Cell Signaling Technology, Cat. #8553S). Protein samples were prepared with NuPAGE LDS Sample Buffer (Thermo Fischer Scientific, Cat. #NP0007), NuPage Sample Reducing Agent (Thermo Fischer Scientific, Cat. #NP0004) or 2X Laemmli buffer and heated at 70 °C or 94 °C for 10 minutes. Samples were resolved by SDS-PAGE, transferred to nitrocellulose (Bio-Rad) and blocked in 5% milk/TBST or 5% milk/PBST for 1 h at room temperature. Membranes were incubated overnight with the respective primary antibodies at 4 °C in 5% milk/PBST or in 5% BSA/TBST, washed with TBST or PBST, incubated with HRP-conjugated secondary antibodies and developed using SuperSignal^TM^ West Dura, the SuperSignal^TM^ West Pico PLUS, or the SuperSignal^TM^ West Femto kits (ThermoFisher Scientific, Cat. 34076, Cat. 34580 and Cat. 34096).

**Supplementary Table 2:**
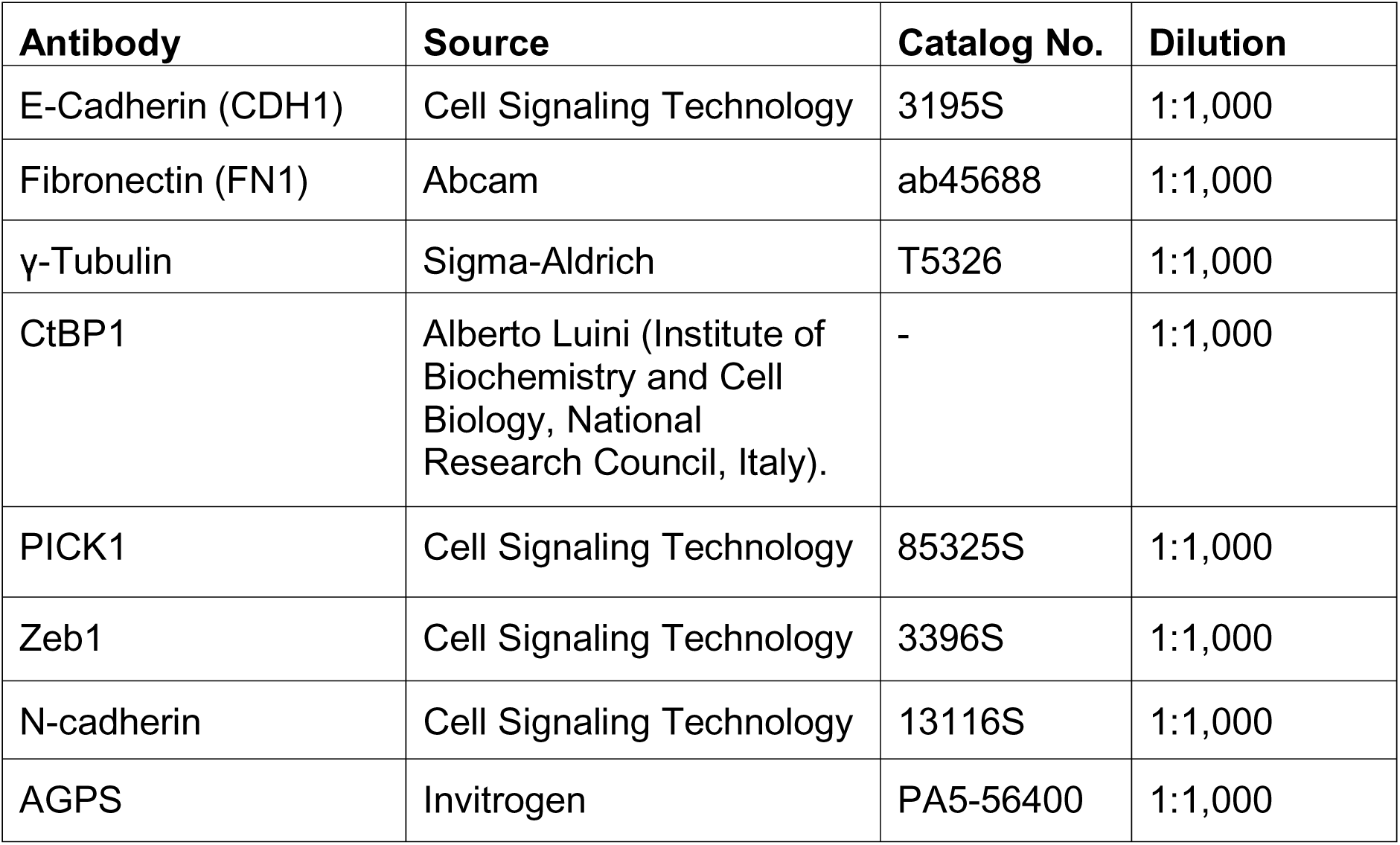
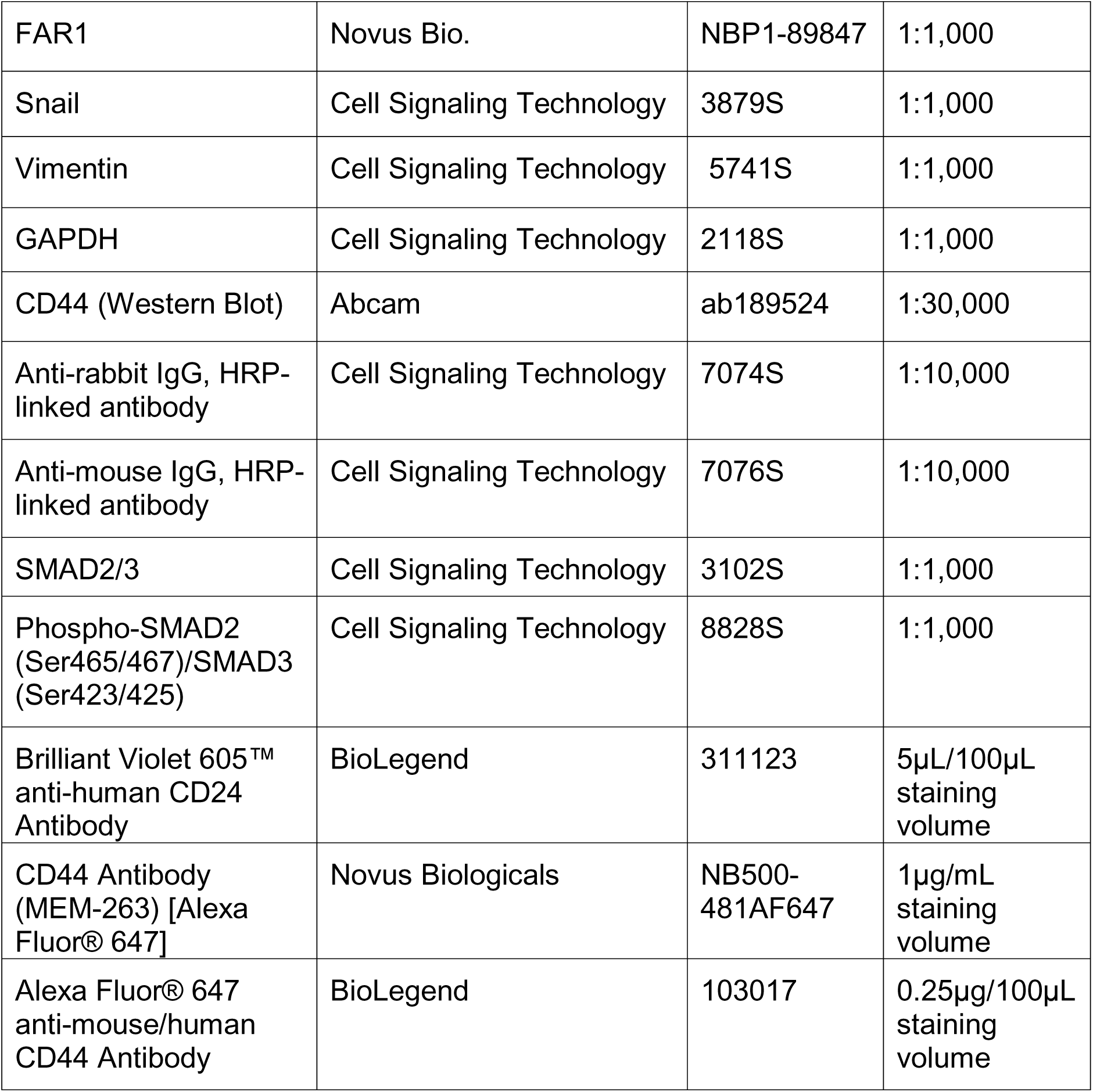
Antibody information. The table below indicates the antibodies used for western blotting and flow cytometry analyses.

| Antibody | Source | Catalog No. | Dilution |
| --- | --- | --- | --- |
| E-Cadherin (CDH1) | Cell Signaling Technology | 3195S | 1:1,000 |
| Fibronectin (FN1) | Abcam | ab45688 | 1:1,000 |
| γ-Tubulin | Sigma-Aldrich | T5326 | 1:1,000 |
| CtBP1 | Alberto Luini (Institute of Biochemistry and Cell Biology, National Research Council, Italy). | - | 1:1,000 |
| PICK1 | Cell Signaling Technology | 85325S | 1:1,000 |
| Zeb1 | Cell Signaling Technology | 3396S | 1:1,000 |
| N-cadherin | Cell Signaling Technology | 13116S | 1:1,000 |
| AGPS | Invitrogen | PA5-56400 | 1:1,000 |
| FAR1 | Novus Bio. | NBP1-89847 | 1:1,000 |
| Snail | Cell Signaling Technology | 3879S | 1:1,000 |
| Vimentin | Cell Signaling Technology | 5741S | 1:1,000 |
| GAPDH | Cell Signaling Technology | 2118S | 1:1,000 |
| CD44 (Western Blot) | Abcam | ab189524 | 1:30,000 |
| Anti-rabbit IgG, HRP-linked antibody | Cell Signaling Technology | 7074S | 1:10,000 |
| Anti-mouse IgG, HRP-linked antibody | Cell Signaling Technology | 7076S | 1:10,000 |
| SMAD2/3 | Cell Signaling Technology | 3102S | 1:1,000 |
| Phospho-SMAD2 (Ser465/467)/SMAD3 (Ser423/425) | Cell Signaling Technology | 8828S | 1:1,000 |
| Brilliant Violet 605™ anti-human CD24 Antibody | BioLegend | 311123 | 5µL/100µL staining volume |
| CD44 Antibody (MEM-263) [Alexa Fluor® 647] | Novus Biologicals | NB500-481AF647 | 1µg/mL staining volume |
| Alexa Fluor® 647 anti-mouse/human CD44 Antibody | BioLegend | 103017 | 0.25µg/100µL staining volume |

### Cell viability assay

Cells were seeded in 96-well black clear bottom plates (Corning) at 2000 or 3000 cells (PyMT-1099 +/− TGF-β) and 6000 cells (OVCAR8) per well. Approximately, 12-16 h post-seeding, cells were treated with various drug concentrations using an HP D300e Digital Dispenser unless stated otherwise. Cell viability was assessed at 72 h posttreatment by performing CellTiter-Glo Luminescent Cell Viability Assays (Promega) according to the manufacturer’s instructions. Relative viability was calculated by normalizing to untreated controls unless stated otherwise. Non-linear regression models (four parameter, variable slope) were applied to generate the regression fit curves using GraphPad Prism version 10.4.1. Drug compounds were purchased as indicated: RSL3 (Selleck Chem), ML210 (Sigma Aldrich), and Liproxstatin-1 (Fisher Scientific). For experiments involving ferric ammonium citrate (FAC), FAC (Sigma) was prepared fresh in sterile 1× PBS and manually added directly to cell culture media at the indicated concentrations at the time of seeding into 96-well plates. Unless stated otherwise, cells were pretreated with FAC for 24 h prior to ML210 treatment.

### Inductively coupled plasma mass spectrometry (ICP-MS)

For select experiments, cells were treated for 24 h with hyaluronic acid (Carbosynth, FH45321, 600-1000 kDa, 1 mg/mL) or hyaluronidase (HD, Sigma-Aldrich, H3884, 0.1 mg/mL) as indicated. Glass vials equipped with Teflon septa were cleaned with nitric acid 65% (VWR, Suprapur, 1.00441.0250), washed with ultrapure water (Sigma-Aldrich, 1012620500) and dried. Cells were harvested and washed twice with 1× PBS. Cells were then counted using an automated cell counter (Entek) and transferred in 200µL 1× PBS to the cleaned glass vials. The same volume of PBS was transferred into separate vials for the background subtraction, at least in duplicate per experiment. For tumor samples, small pieces of the tumors were added into pre-weighed cleaned glass vials. Samples were lyophilized using a freeze dryer (CHRIST, 22080). Glass vials with lyophilized tumor samples were weighed to determine the dry weight for normalization. Samples were subsequently mixed with nitric acid 65% and heated at 80°C overnight. Samples were diluted with ultrapure water to a final concentration of 0.475 N nitric acid and transferred to metal-free centrifuge vials (VWR, 89049-172) for subsequent ICP-MS analyses. Amounts of metals were measured using an Agilent 7900 ICP-QMS in low-resolution mode, taking natural isotope distribution into account. Sample introduction was achieved with a micro-nebulizer (MicroMist, 0.2 mL/min) through a Scott spray chamber. Isotopes were measured using a collision-reaction interface with helium gas (5 mL/min) to remove polyatomic interferences. Scandium and indium internal standards were injected after inline mixing with the samples to control the absence of signal drift and matrix effects. A mix of certified standards was measured at concentrations spanning those of the samples to convert count measurements to concentrations in the solution. Values were normalized against cell number or dry weight.

### Iron measurements using HRhoNox-M

The lysosome-specific fluorescent Fe(II) probe HRhoNox-M was synthesized in 3 steps according to a previously published procedure^36^. ^1^H NMR (300 MHz, CDCl_3_) δ 7.93 (1H, d, *J* = 2.0 Hz), 7.45 (1H, dd, *J* = 8.5 Hz, 2.0 Hz), 7.40–7.30 (3H, m), 7.05 (1H, d, *J* = 8.5 Hz), 6.90 (1H, d, *J* = 7.0 Hz), 6.80 (1H, d, *J* = 8.0 Hz), 6.50–6.44 (2H, m), 5.28–5.35 (2H, m), 3.62 (6H, m), 2.97 (6H, s). MS (ESI) m/z: calcd. for C_24_H_25_N_2_O_3_ [M+H]^+^ 389.19, found: 389.35. Cells were incubated with 1 µM HRhoNox-M for 1 h or lysotracker deep red (Thermo Fisher Scientific L12492) according to the manufacturer’s instructions for 1 h. Cells were then washed twice with ice-cold PBS and suspended in incubation buffer prior to being analyzed by flow cytometry. For each condition, at least 10,000 cells were counted. Data were recorded on a BD Accuri C6 (BD Biosciences) and processed using Cell Quest (BD Biosciences) and FlowJo (FLOWJO, LLC). The signal for HRhoNox-M was normalized against the signal of lysotracker of cells treated in parallel.

### Endocytosis experiments

Antibody against EEA1 was purchased from BD Biosciences (Catalog no. 610456). Conjugated transferrin (mouse)-Alexa546, dextran-Alexa555, and EGF-Alexa555 were purchased from Invitrogen. Cy3-conjugated hyaluronate was synthesized in-house. Cy2- or Cy3-conjugated donkey antibodies against mouse IgG were purchased from Jackson ImmunoResearch. For receptor-mediated endocytosis, cells were washed with serum-free medium and then incubated in this medium with Cy3-conjugated hyaluronate (0.1 mg/ml), Alexa 555-conjugated EGF (either 2 ng/ml or 200 ng/ml), or Alexa 546-conjugated transferrin (5 μg/ml) for 1 h at 4° C. Cells were then washed to clear unbound ligand, and shifted to 37 °C for times indicated in the figures. Cells were stained for EEA1, followed by confocal microscopy to assess the arrival of ligand to the early endosome. To assess fluid-phase uptake, Alexa 555-conjugated dextran (0.2 mg/ml) was added to complete medium and cells were incubated 37 °C for times indicated in the figures. Cells were then stained for EEA1, followed by confocal microscopy to assess the arrival of this probe to the early endosome. In select assays cells were pretreated with 20μM C(22:6) BSA conjugate for 16-18 h.

### Confocal microscopy

Colocalization studies were performed with the Zeiss Axio Observer Z1 Inverted Microscope having a Plan-Apochromat 63x objective, the Zeiss LSM 800 with Airyscan confocal package with Zeiss URGB (488- and 561-nm) laser lines, and Zen 2.3 blue edition confocal acquisition software. For quantification of colocalization, ten fields of cells were examined, with each field typically containing about 5 cells. Images were imported into the NIH ImageJ v.1.50e software, and then analyzed through a plugin software (https://imagej.net/Coloc_2). Under the ‘image’ tab, the ‘split channels’ option was selected. Under the ‘plugins’ tab, ‘colocalization analysis’ option was selected, and within this option, the ‘colocalization threshold’ option was selected. Menders Coefficient was used for colocalization analysis. Colocalization values were calculated by the software and expressed as the fraction of protein of interest colocalized with EEA1. Colocalization experiments were performed at least twice, and representative data is shown.

### Mammosphere Assay

Mammospheres were generated in accordance to a previously described protocol with minor modifications^114^. Briefly, single cell suspensions containing 1,200 WT or AGPS KO pB3 cells were plated in DMEM/F12 supplemented with 2nM L-glutamine, 20ng/mL recombinant human epidermal growth factor (EGF; Peprotech Cat. No. AF-100-15), and B27 supplement (Gibco, Cat. No. 17504-044) in individual wells of an ultra-low attachment 6 well plate (Corning Cat. No. 3471). After 4 days, the total number of spheres greater than 40 µm were quantified before dissociation in trypsin at 37°C. Dissociated cells were then passed through a 40 µm cell strainer to yield a single cell suspension, before replating 1,200 cells. Sphere formation efficiency ((# of mammospheres per well) / (# of cells seeded per well) x 100) was calculated over 3 passages. Data shown is representative of 3 independent biological replicates.

### Synthesis of HA-Cy3 probe

Hyaluronic acid (HA, 2 mg, Sigma 75044, Lot #BCBM2884) was dissolved in a 1:1 solution of dimethylsulfoxide (DMSO) and water (0.4 mL) for a stock concentration of 5 mg/mL. The polymer was sonicated under heating to ensure full solubilization. The HA solution was then diluted into HEPES (50 mM final HEPES concentration for a total reaction volume of 2 mL once all components are combined). Sulfo-Cyanine3 amine (2.36 mg, Lumiprobe) was separately dissolved in DMSO (0.236 mL) for a stock concentration of 10 mg/mL. N-(3-Dimethylaminopropyl)-N′-ethylcarbodiimide hydrochloride (EDC, 0.253 mg, Sigma) was separately dissolved in 50 mM HEPES (0.051 mL) for a stock concentration of 5 mg/mL. The HA and EDC solutions were then combined under stirring, followed by addition of the dye solution. The reaction was stirred, protected from light, at room temperature for 12 h. Following, unreacted dye was removed via Amicon Ultra-0.5 Centrifugal Filters (Millipore Sigma). Manufacturer guidelines were followed to select purification spin speeds and times: 14000 xg, 15 min per wash step (water) until washes were clear and colorless. The purified HA-Cy3 probe was stored in water at 4 °C until used.

### Preparation of liposomes

Ether lipid liposomes were prepared as previously described^22^. PE (18:1p_20:4) (Catalog no. 852804), PE (18:1p_18:1) (Catalog no. 852758), PC (18:1p_20:4) (Catalog no. 852469) and PE (18:0_20:4) (Catalog no. 850804) were purchased from Avanti Polar Lipids Inc.

**Supplementary Table 3:** Characterization of liposomes. Table showing the hydrodynamic diameter and polydispersity of liposomes used in this study. The average and standard deviation of three technical repeats is provided. A Malvern ZS90 Particle Analyzer was used for size measurements reported.

| Liposome | Hydrodynamic diameter (nm) | Polydispersity index |
| --- | --- | --- |
| PE (18:1p_20:4) | 174.8 ± 2.25 | 0.273 ± 0.01 |
| PE (18:1p_18:1) | 128.7 ± 2.07 | 0.287 ± 0.03 |
| PC (18:1p_20:4) | 146.8 ± 1.52 | 0.21 ± 0.002 |
| PE (18:0_20:4) | 182.0 ± 0.13 | 0.181 ± 0.02 |

### Liposome reconstitution assays

Adherent cells were treated with liposomes 16-18 h prior to performing respective membrane tension or endocytosis assays. Lipid liposomes were added directly to the culture medium for a final concentration of 20 µM. Cells were switched from liposome-containing media to extracellular imaging buffer (HEPES buffer with dextrose, NaCl, KCl, MgCl2, CaCl2) during membrane tension experiments.

### Preparation of BSA conjugated DHA C(22:6)

In brief, BSA (FA-free; Fisher Scientific BP9704100) was dissolved in ddH2O to give a 1 mM solution. Exactly 25 mg of docosahexaenoic acid (DHA, C22:6; Sigma Aldrich D2534) was added to 38.05 mL of 1 mM BSA solution to yield a 2:1 DHA:BSA ratio in a glass vial. The vial was then purged with nitrogen and stirred at room temperature until dissolved. The conjugate was then filter sterilized and concentration verified via BCA assay. Aliquots were then stored at -80C until use.

### Membrane tension

Tether pulling experiments were performed on a home-built optical trap, following principles described elsewhere^74, 115^. Briefly, 4 µm anti-Digoxigenin-coated polystyrene beads (Spherotech) were trapped with a 1064 nm, Ytterbium laser (IPG Photonics) focused through a 60x 1.2 NA objective (Olympus). Forces on the beads were measured by the deflection of backscattered trapping laser light onto a lateral effect position sensor (Thorlabs) and calibrated using the viscous drag method^116^. To measure tether radii (R), cell lines were transiently transfected with a membrane-targeted fluorescent protein (glycosylphosphatidylinositol-anchored eGFP, Addgene #32601) using a TransIT-X2 transfection kit (Mirus). Tether radius was obtained by comparing tether fluorescence to fluorescence counts from a known area of the parent cell membrane, as described^59^. Tether force ( and fluorescence measurements were performed simultaneously. Membrane tension was calculated using the following equation:

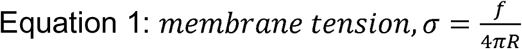

### Miscibility transition temperatures (T_misc_) measurements

Miscibility transition temperatures (T_misc_) measurements were performed as previously reported^106, 117^. Briefly, cells were washed in PBS, and cell membranes were labeled with 5 µg/ml fluorescent disordered/nonraft phase marker FAST DiO (Thermo Fisher Scientific) for 10 min on ice. Cells were then washed twice in GPMV buffer (10 mM HEPES, 150 mM NaCl, 2 mM CaCl2, pH 7.4), and then incubated with GPMV buffer supplemented with 25 mM paraformaldehyde (PFA) and 2 mM dithiothreitol (DTT) for 1 h at 37 °C. Vesicles were imaged at 40× on an inverted epifluorescence microscope (Leica DMi8) under temperature-controlled conditions using a microscope stage equipped with a Peltier element (Warner Instruments). GPMVs were imaged from 4° C28 °C, counting phase-separated and uniform vesicles at each temperature. For each temperature, 25-50 vesicles were counted, and the percent of phase-separated vesicles were calculated, plotted versus temperature, and a fitted to a sigmoidal curve to determine the temperature at which 50% of the vesicles were phase-separated (T_misc_).

### C-laurdan spectral imaging

C-Laurdan imaging was performed as previously described^80, 82, 106, 117, 118^. Briefly, cells were washed with PBS and stained with 10 µg/mL C-Laurdan for 10 min on ice, then imaged using confocal microscopy on a Leica SP8 with spectral imaging at 60× (water immersion, NA= X) and excitation at 405 nm. The emission was collected as two images: 420–460 nm and 470–510 nm. MATLAB (MathWorks, Natick, MA) was used to calculate the two-dimensional (2D) GP map, where GP for each pixel was calculated as previously described ^118^. Briefly, each image was background subtracted and thresholded to keep only pixels with intensities greater than 3 standard deviations of the background value in both channels. The GP image was calculated for each pixel using Equation 2, where G is the G-factor. The G-factor was determined before each experiment using the protocol in^119^. GP maps (pixels represented by GP value rather than intensity) were imported into ImageJ. To calculate the average PM GP, line scans drawn across individual cells. PM GP values were taken as peak GP values from the periphery of the cell, whereas internal membranes were calculated as the average of all values outside the PM peak. The average GP of the internal membranes was calculated by determining the average GP of all pixels in a mask drawn on each cell just inside of the PM.

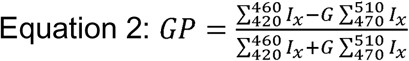

### Extravasation assay

#### Cells and reagents

Immortalized human umbilical vein endothelial cells (ECs) expressing BFP^95^ were cultured in VascuLife VEGF Endothelial Medium (Lifeline Cell Technology). Normal human lung fibroblasts (FBs) (Lonza, P7) were cultured in FibroLife S2 Fibroblast Medium (Lifeline Cell Technology).

#### Microfluidic device

3D cell culture chips (AIM Biotech) were used to generate in vitro microvascular networks (MVNs). The AIM chip body was made of cyclic olefin polymer (COP) with a type of gas-permeable plastic serving as the bottom film. AIM Biotech chips contained three parallel channels: a central gel channel flanked by two media channels.

Microposts separated fluidic channels and serve to confine the liquid gelling solution in the central channel by surface tension before polymerization. The gel channel was 1.3 mm wide and 0.25 mm tall, the gap between microposts was 0.1 mm, and the width of media channels was 0.5 mm.

#### Microvascular network formation

To generate perfusable MVNs, ECs and FBs were seeded into the microfluidic chip using a two-step method^120^. Briefly, ECs and FBs were concentrated in VascuLife containing thrombin (4 U/mL). For the first step seeding, the outer layer EC solution was made with a final concentration of 10 ×10^6^/mL. After mixed with fibrinogen (3 mg/mL final concentration) at a 1:1 ratio, the outer layer EC solution was pipetted into the gel inlet, immediately followed by aspirating from the gel outlet, leaving only residual solution around the microposts. For the second step, another solution with final concentrations of 5 ×10^6^/mL ECs and 1.5 ×10^6^/mL fibroblasts were similarly mixed with fibrinogen and then pipetted into the same chip through the gel outlet. The device was placed upside down to polymerize in a humidified enclosure and allowed to polymerize at 37 °C for 15 min in a 5% CO_2_ incubator. Next, VascuLife culture medium was added to the media channels and changed daily in the device. After 7 days, MVNs were ready for further experiments.

#### Tumor cell perfusion in MVNs

PyMT-1099 or pB3 cell line derivatives expressing pCDH-EF1-Luc2-P2AtdTomato (Plasmid #72486, Addgene) were resuspended at a concentration of 1×10^6^/mL in culture medium. To perfuse these tumor cells into in vitro MVNs, the culture medium in one media channel was aspirated, followed by injection of a 20 µL tumor cell suspension in the MVNs and repeated twice. Microfluidic devices were then placed at 37 °C for 15 min in a 5% CO_2_ incubator for 15 min. After that, the tumor cell medium was aspirated from the media channels to remove the unattached cells, and Vasculife was replenished. Devices were then placed back to the incubator. 24 h later, devices were fixed, washed, and imaged using an Olympus FLUOVIEW FV1200 confocal laser scanning microscope with a 10× objective and an additional 2× zoom-in function. Zstack images were acquired with a 5 µm step size. All images shown are collapsed Zstacks, displayed using range-adjusted Imaris software, unless otherwise specified. Extravasation percentage was calculated by dividing the cell number of extravasated tumor cells with the total number of tumor cells in the same imaging region of interest.

### CLIC/GEEC and micropinocytosis inhibition

7-keto-cholesterol was obtained from Cayman Chemical (Cat. 16339). Cytochalasin D was obtained from Sigma Aldrich (C8273). pB3 cells were pre-treated for 30 minutes with either of the inhibitors before assessing endocytic rate.

### RNA interference

In brief, cells were seeded in 6-well plates at the density of 2×10^5^ cells/well and transfected 24hr later with the specified siRNA using jetPRIME (Polyplus, 114-15), or PepMute™ (SignaGen, Cat. SL100566) according to the manufacturer’s protocol with 100 nM siRNA. The medium was replaced after 6 h. Analysis was performed 72 h after transfection. Suitable small interfering RNAs were designed by Dharmacon for specific down-regulation of PICK1 (5’-cuuagacuaugacaucgaa-3) and CtBP1 (5’ugucucaucugcuugacagu-3).

### Flow Cytometry

MCF7 cells were washed with ice-cold PBS. For antibody staining, cells were incubated with Fc block (Human TruStain FcX, Biolegend, 422302, 1:20) for 15 min, then incubated with the following antibodies: CD24-BV605 (Biolegend, 311124) and CD44AF647 (Novus Biologicals, NB500-481AF647) in ice cold 10% FBS, PBS, 2 mM EDTA for 20 min at 4 °C and then washed with PBS and resuspended in 10% FBS, PBS, 2 mM EDTA before analysis using a Attune flow cytometer. Data were analyzed with FlowJo software v. 10.10.0.

### Histology

Harvested tissues were fixed by incubating with 10% neural-buffered formalin (VWR Scientific) at 4°C for 16–18 h. Fixed samples were then transferred to 70% ethanol and submitted to Hope Babette Tang Histology Facility at the Koch Institute at MIT for paraffin-embedding and H&E staining. Metastatic burden was quantified using QuPath software^121^ and Image J^122^.

### Statistical Analysis

For statistical analyses, Mann-Whitney U test, two-tailed t-test, and one way ANOVA with either Dunnett, Tukey, or Holm-Šídák multiple test corrections were performed using GraphPad Prism Version 10.4.1. Permutation ANOVA was performed in R (v4.4.2) using the *coin* package with 10,000 permutations^123^. Planned contrasts were tested with one-sided Welch’s *t*-tests (t.test, base R), and exploratory pairwise comparisons were adjusted for multiple testing using Holm’s method (p.adjust).

#### Data Availability

Lipidomic analysis of ether-lipid-deficient cells has been deposited in Zenodo (<u>10.5281/zenodo.17808937</u>). All other data is provided within source data.

Correspondence and requests for materials should be addressed to: Raphaël Rodriguez, PhD, Robert A. Weinberg, PhD and Whitney S. Henry, PhD.

## ACKNOWLEDGMENTS

We thank all members of the Weinberg, Farese and Walther labs for insightful discussions and reagents. We are grateful to Brent Stockwell, Laurie Boyer, Edward Lyman, and Fred Heberle for helpful discussions. We acknowledge technical support from Caroline A. Lewis, Zon W. Lai and Marina Plays, the CurieCoreTech Metabolomics and Lipidomics Technology Platform at the Institut Curie, the following facilities at the Whitehead Institute: Metabolite Profiling Core, W.M. Keck Microscopy Core, Flow Cytometry Core, as well as the Koch Institute’s Robert A. Swanson (1969) Biotechnology Center, specifically the MIT Koch Institute Animal Imaging and PreClinical Testing Core Facility and Hope Babette Tang (1983) Histology Core Facility. We also thank Naviya Kapadia for assistance with proofreading and providing helpful comments on the manuscript and figures.

This work was funded in part by the National Institutes of Health: Koch Institute Support (core) Grant P30CA014051 from the National Cancer Institute (WSH), National Institute Of General Medical Sciences R37GM058615 (VWH), the National Cancer Institute K99CA255844/R00CA255844 (NB), and the National Cancer Institute (RDK). Additional funding support was received from MIT Stem Cell (RAW), Brendan Bradley Gift (RAW), Nile Albright Research Foundation (RAW), Samuel Waxman Cancer Research Foundation (RAW), Virginia and D.K. Ludwig Fund for Cancer Research Center (RAW), Marble Center for Cancer Nanomedicine (PTH), Jane Coffin Childs Memorial Fund (WSH), Ludwig Center at MIT’s Koch Institute for Integrative Cancer Research (WSH), Emerald Foundation, Inc. (WSH), National Institute Of General Medical Sciences 1R21CA300756 (KRL), National Institute Of General Medical Sciences R35GM134949 (IL), National Institute Of General Medical Sciences T32GM136540 (RPM & JS) and the New Horizon UROP Fund/MIT (VVP). R.R. is supported by Ligue Contre le Cancer (Equipe Labellisée), Fondation Charles DefforeyInstitut de France, Fondation Bettencourt Schueller, Institut National du Cancer, Agence Nationale de la Recherche and the European Research Council under the European Union’s Horizon 2020 research and innovation program (grant agreement no. 647973). R.R. also received support from INCa 2022-1-PL BIO-05-ICR-1 and INCa_16712.

## AUTHOR CONTRIBUTIONS

W.S.H. and R.R. conceived the project. W.S.H, R.P.M. and S.M. designed and performed experiments with input from R.A.W. and R.R. S.M. conducted iron measurement studies. S.M. and J.L.S. performed oxidized lipidomics analysis. J.Y. performed endocytosis assays, with critical insight from V.W.H. Membrane tension studies were conducted by S.I., with support from A.E.C. on data analysis. K.R.L. performed membrane tension experiments and together with I.L. provided key insights on interpretation. W.S.H., S.D. and F.R. conducted animal experiments. Z.W. performed extravasation assays with support from R.D.K. A.D. and C.B.C provided significant evaluation and interpretation of lipidomics data. Liposomal nanoparticles and HA-Cy3 probes were synthesized by N.B. and A.V. G.W.B. and P.T. provided support for data analysis. K.J.S., V.V.P., J.S., D.H., F.S., L.S., N.S. and E.N.E. contributed to data acquisition and analysis. P.T.H. provided helpful discussion and scientific input. W.S.H., and R.P.M. wrote the manuscript. W.S.H., R.A.W., and R.R. edited the manuscript with input from all authors.

## FIGURE LEGENDS

**Supplementary Fig. 1.**
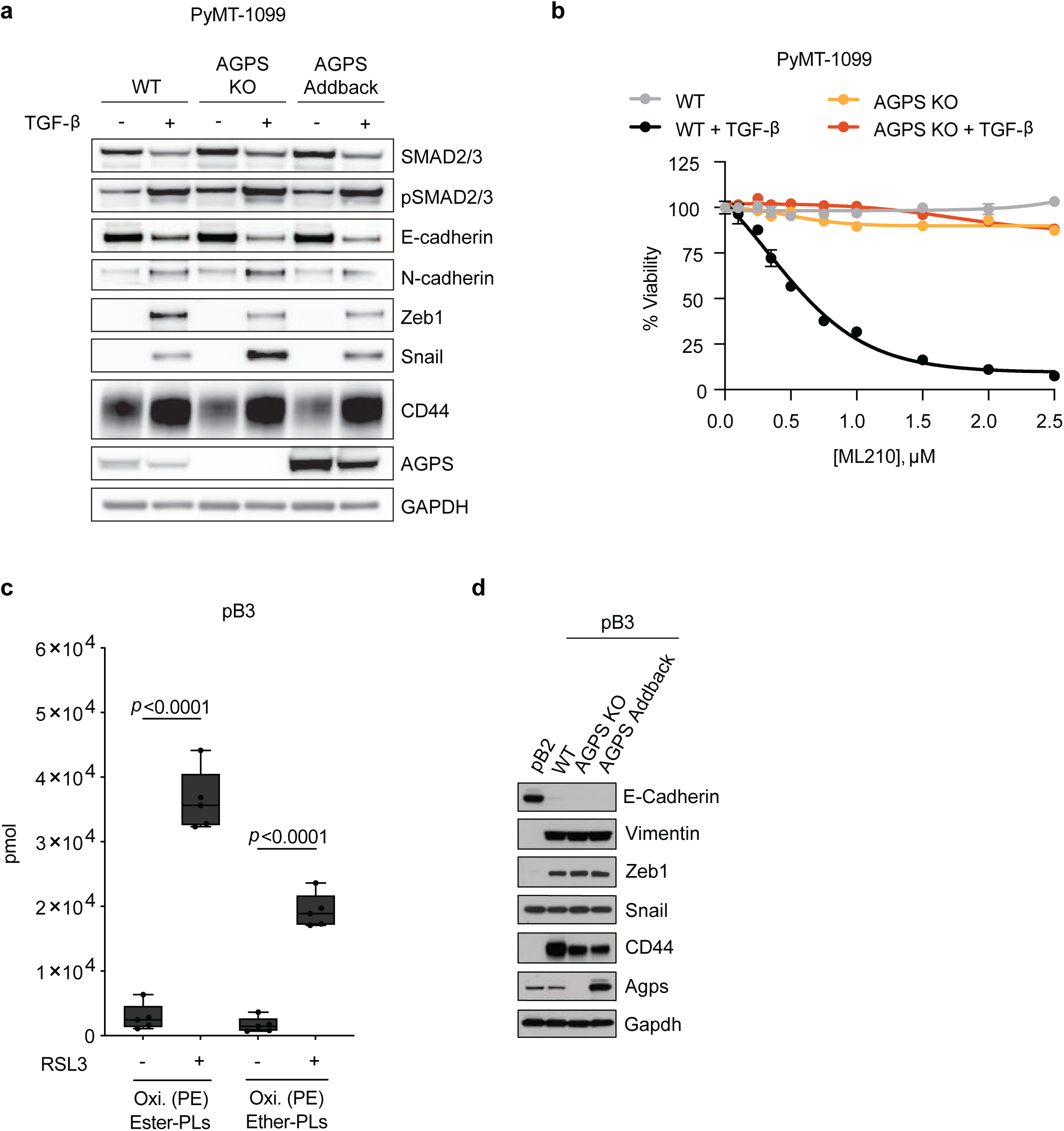
a. Immunoblot analysis for EMT markers and TGF-β signaling in PyMT-1099 WT or AGPS KO cells. Where indicated cells were treated with TGF-β (2 ng/ml) for 10 d. b. Cell viability following treatment with the GPX4 inhibitor ML210 for 72 h (n=3). PyMT-1099 WT or AGPS KO cells were pretreated with TGF-β (2 ng/ml) for 10 d prior to assay. Graph is representative of three independent biological replicates. Data shown as mean +/− SEM. c. Amount in pmol of oxidized phosphatidylethanolamine (Oxi. PE) ether and ester phospholipids in pB3 cells treated with RSL3 or vehicle control for 24 hours (n=5). 100,000 cells were used for lipid extraction in each condition. Data shown as mean +/− SEM. Statistical significance was calculated using unpaired, two-tailed t-test. d. Immunoblot analysis for AGPS expression in mesenchymal-enriched pB3 WT, AGPS KO, and AGPS addback cells. pB2 cells served as a control for expression of epithelial-like markers. For panel a: PyMT-1099 WT and AGPS KO cells were transduced with the respective vector control plasmids. For panel d: pB3 WT and AGPS KO cells were transduced with the respective vector control plasmids.

**Supplementary Fig. 2.**
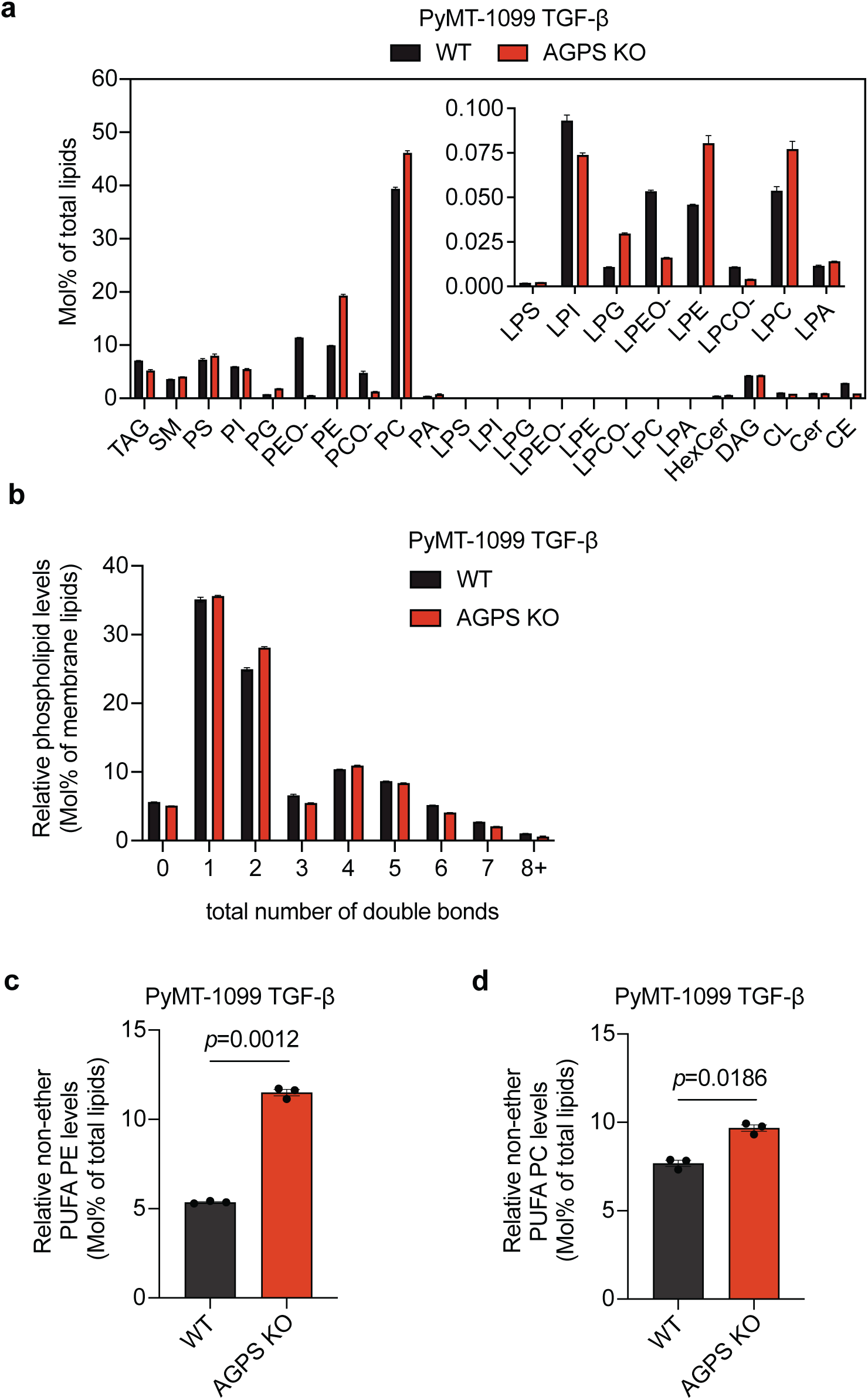
a. Relative abundance of lipid classes in PyMT-1099 WT and AGPS KO cells treated with TGF-β (2 ng/ml, 10 d). Lipid classes include triacylglycerol (TAG), sphingomyelin (SM), phosphatidylserine (PS), phosphatidylinositol (PI), phosphatidylglycerol (PG), phosphatidylethanolamine (PE), ether phosphatidylethanolamine (PE O-), phosphatidylcholine (PC), ether phosphatidylcholine (PC O-), phosphatidic acid (PA), lysophosphatidylserine (LPS), lysophosphatidylinositol (LPI), lysophosphatidylglycerol (LPG), lysophosphatidylethanolamine (LPE), lysophosphatidylcholine (LPC), ether lysophosphatidylethanolamine (LPEO-), ether lysophosphatidylcholine (LPCO-), lysophosphatidic acid (LPA), hexosylceramide (HexCer), diacylglycerol (DAG), cardiolipin (CL), ceramide (Cer), and cholesteryl ester (CE). b. Bar plot showing the distribution of saturation levels (total number of double bonds) among membrane lipid species present in PyMT-1099 WT and AGPS KO cells treated with TGF-β (2 ng/ml, 10 d). c. Quantification of the relative abundance of non-ether polyunsaturated phosphatidylethanolamine (PUFA PE) species in PyMT-1099 WT and AGPS KO cells treated with TGF-β (2 ng/ml, 10 d). d. Quantification of the relative abundance of non-ether polyunsaturated phosphatidylcholine (PUFA PC) species in PyMT-1099 WT and AGPS KO cells treated with TGF-β (2 ng/ml, 10 d). e. All data shown as the mean +/− SEM of 3 replicates. Statistical significance was calculated using unpaired, two-tailed t-test.

**Supplementary Fig. 3.**
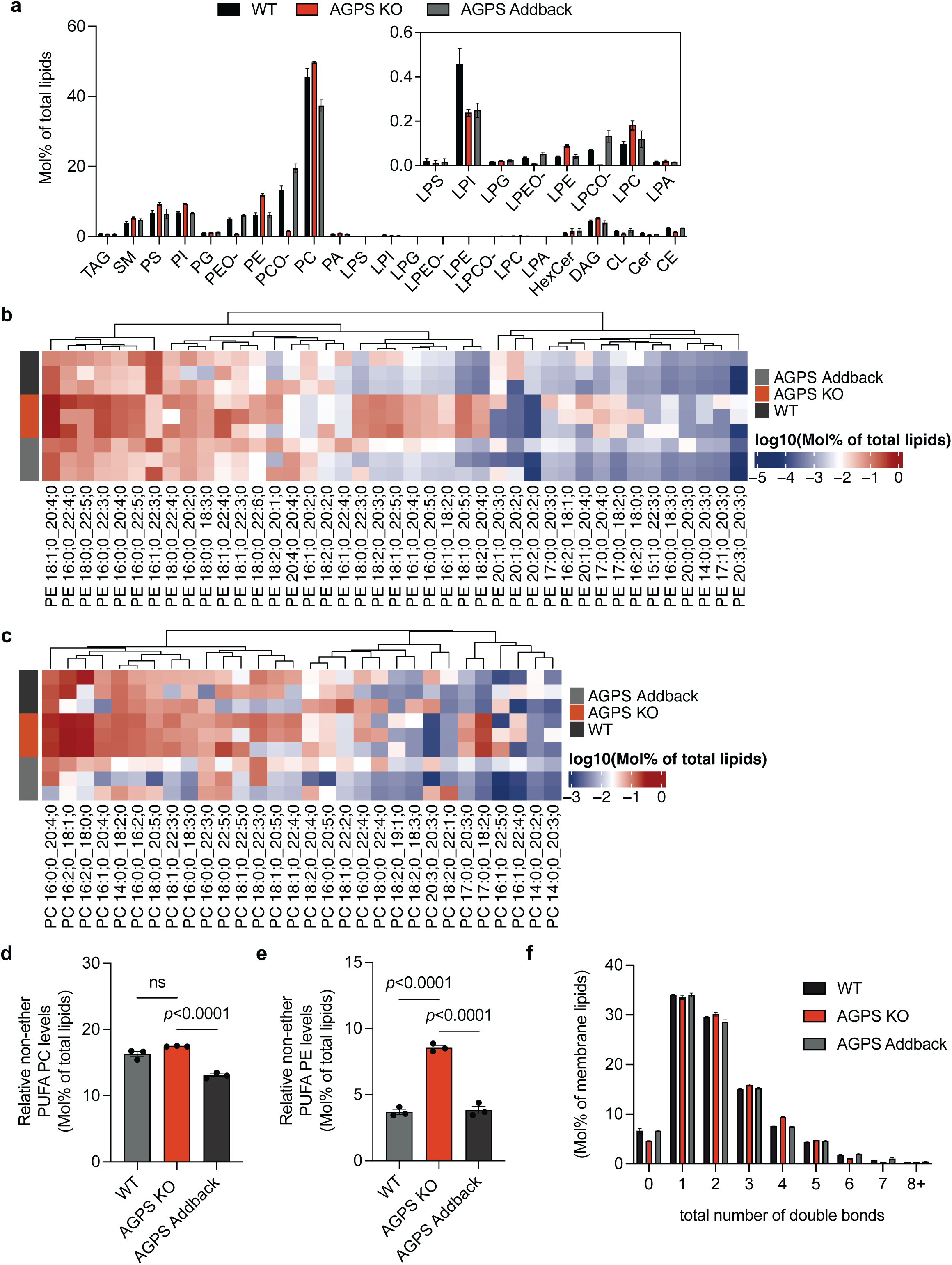
a. Relative abundance of lipid classes in WT, AGPS KO, and AGPS addback pB3 cells. Lipid classes include triacylglycerol (TAG), sphingomyelin (SM), phosphatidylserine (PS), phosphatidylinositol (PI), phosphatidylglycerol (PG), phosphatidylethanolamine (PE), ether phosphatidylethanolamine (PE O-), phosphatidylcholine (PC), ether phosphatidylcholine (PC O-), phosphatidic acid (PA), lysophosphatidylserine (LPS), lysophosphatidylinositol (LPI), lysophosphatidylglycerol (LPG), lysophosphatidylethanolamine (LPE), lysophosphatidylcholine (LPC), ether lysophosphatidylethanolamine (LPEO-), ether lysophosphatidylcholine (LPCO-), lysophosphatidic acid (LPA), hexosylceramide (HexCer), diacylglycerol (DAG), cardiolipin (CL), ceramide (Cer), and cholesteryl ester (CE). b. Hierarchical clustering heatmap of polyunsaturated phosphatidylethanolamine (PUFA PE) species based on abundance (mol% of total lipids) in WT, AGPS KO, and AGPS addback pB3 cells. A sample variance of greater than 10 was used as a filter for representing species within the heatmap. c. Hierarchical clustering heatmap of polyunsaturated phosphatidylcholine (PUFA PC) species based on abundance (mol% of total lipids) in WT, AGPS KO, and AGPS addback pB3 cells. A sample variance of greater than 10 was used as a filter for representing species within the heatmap. d. Bar plot quantifying the relative abundance of non-ether PUFA PC species in WT, AGPS KO, and AGPS addback pB3 cells. e. Bar plot quantifying the relative abundance of non-ether PUFA PE species in WT, AGPS KO, and AGPS addback pB3 cells. f. Bar plot showing the distribution of saturation levels (total number of double bonds) among membrane lipid species present in WT, AGPS KO, and AGPS addback pB3 cells. All samples were analyzed with 3 biological replicates and represented as the mean +/− SEM. Statistical significance was calculated using one-way ANOVA with Tukey’s multiple comparisons test; ns, not significant. pB3 WT and AGPS KO cells were transduced with the respective vector control plasmids.

**Supplementary Fig. 4.**
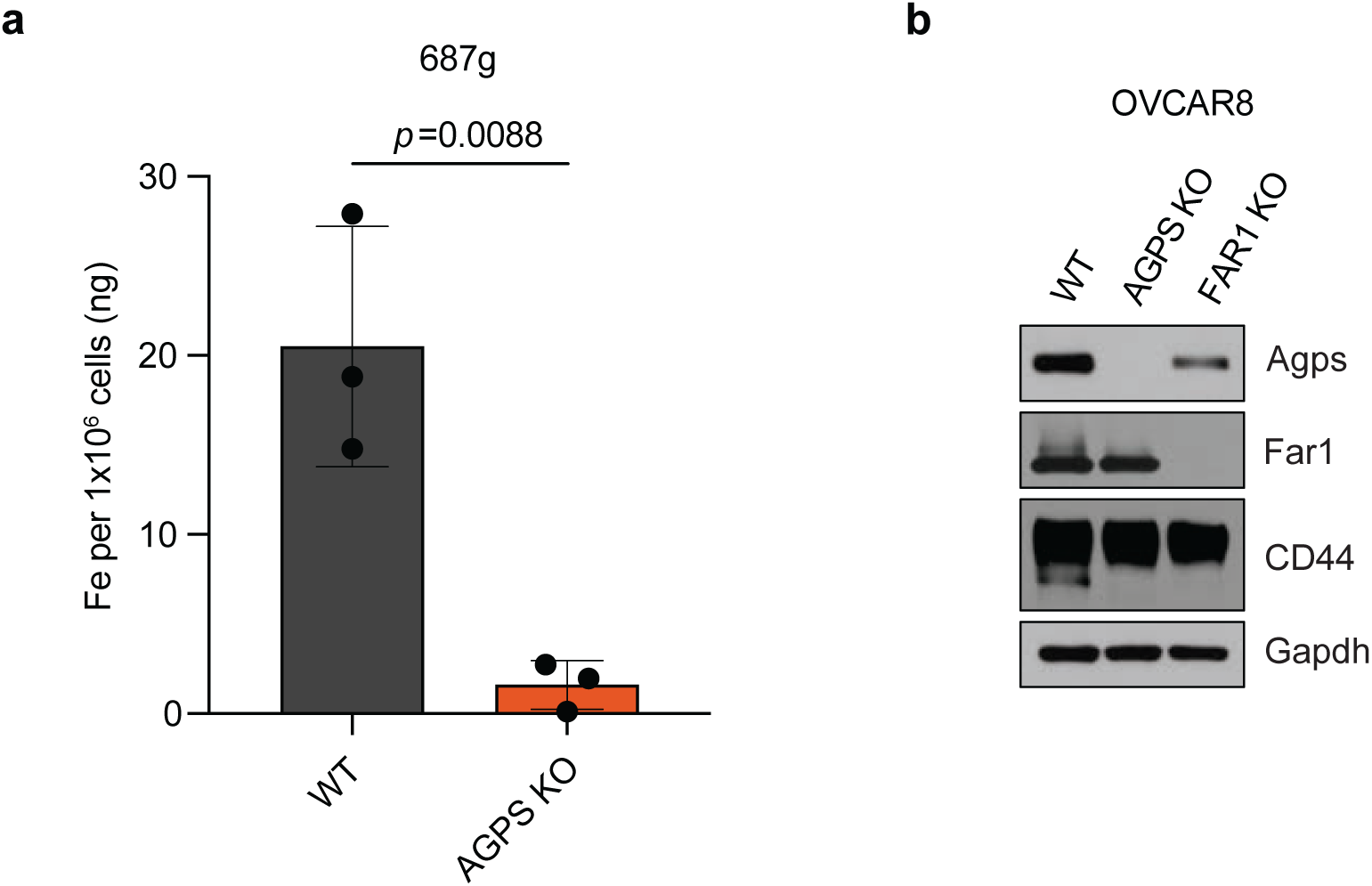
a. ICP-MS of cellular iron in the mesenchymal-enriched 687g WT and AGPS KO murine breast cancer cell line. All shown as the mean +/− SEM of 3 replicates. Statistical significance was calculated using unpaired, two-tailed t-test. b. Immunoblot analysis of OVCAR8 AGPS KO, FAR1 KO or nontargeting sg (WT) cells.

**Supplementary Fig. 5.**
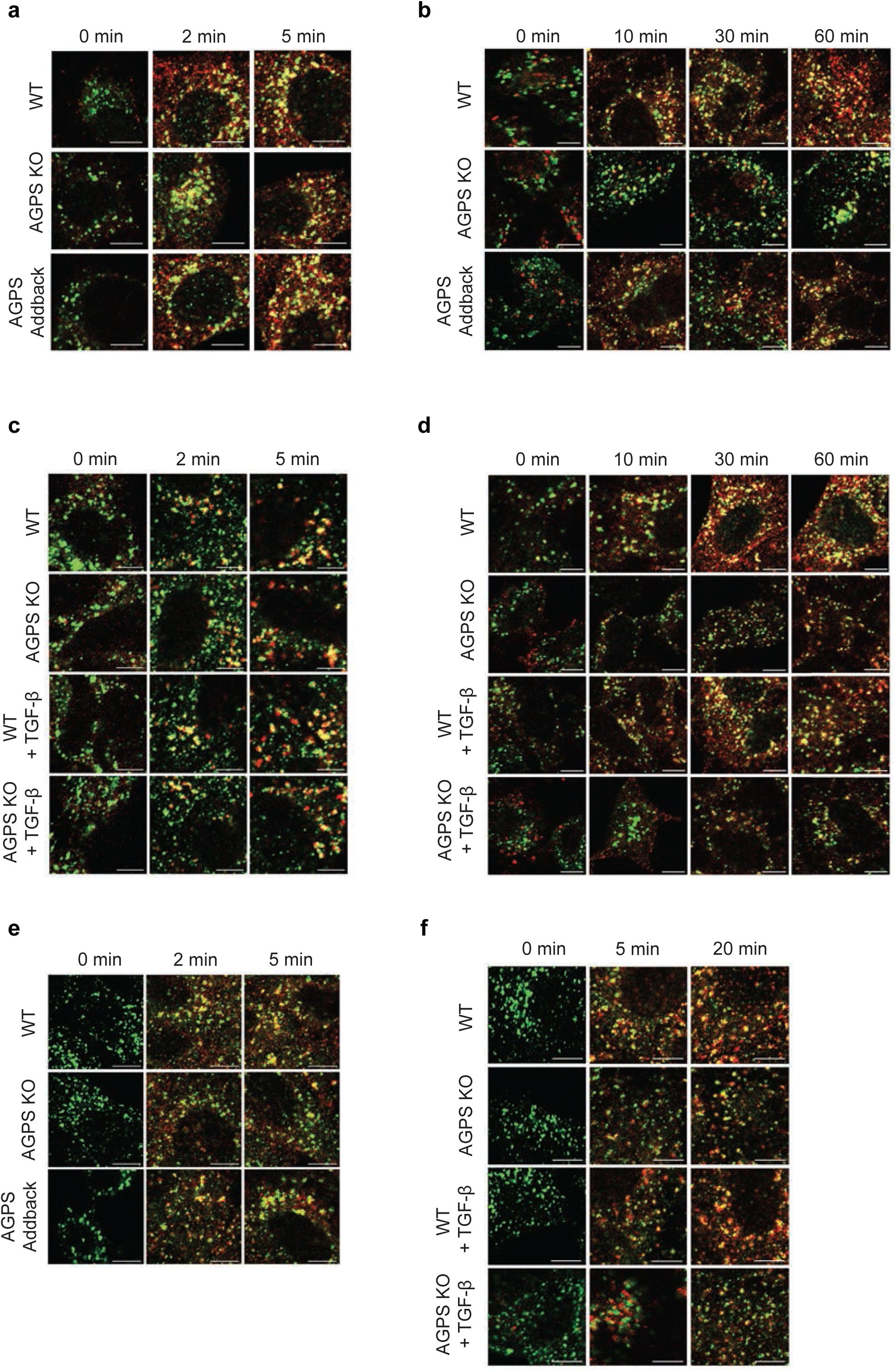
a. Representative confocal images showing internalization of transferrin (red) and colocalization with the early endosome marker EEA1 (green) at 0, 2, and 5 minutes in WT, AGPS KO, and AGPS addback pB3 cells. Scale bar: 10 μm. b. Representative confocal images showing internalization of fluorescently labeled hyaluronate probe (red) and colocalization with EEA1 (green) at 0, 10, 30, and 60 minutes in WT, AGPS KO, and AGPS addback pB3 cells. Scale bar: 10 μm. c. Representative confocal images of transferrin (red) and EEA1 (green) colocalization at 0, 2, and 5 minutes in WT and AGPS KO PyMT-1099 cells +/− TGF-β pretreatment (2 ng/ml; 10 d). Scale bar: 10 μm. d. Representative confocal images showing internalization of fluorescently labeled hyaluronate probe (red) and EEA1 (green) at 0, 10, 30, and 60 minutes in WT and AGPS KO PyMT-1099 cells +/− TGF-β pretreatment (2 ng/ml; 10 d). Scale bar: 10 μm. e. Representative confocal images showing dextran (red) internalization and colocalization with EEA1 (green) at 0, 2, and 5 minutes in WT, AGPS KO, and AGPS addback pB3 cells. Scale bar: 10 μm. f. Representative confocal images of dextran (red) and EEA1 (green) at 0, 5, and 20 minutes in WT and AGPS KO PyMT-1099 cells +/− TGF-β pretreatment (2 ng/ml; 10 d). Scale bar: 10 μm. For panels a, b & e: pB3 WT and AGPS KO cells were transduced with the respective vector control plasmids. Examined n=10 fields of cells per experimental sample and at least two independent replicates were performed.

**Supplementary Fig. 6.**
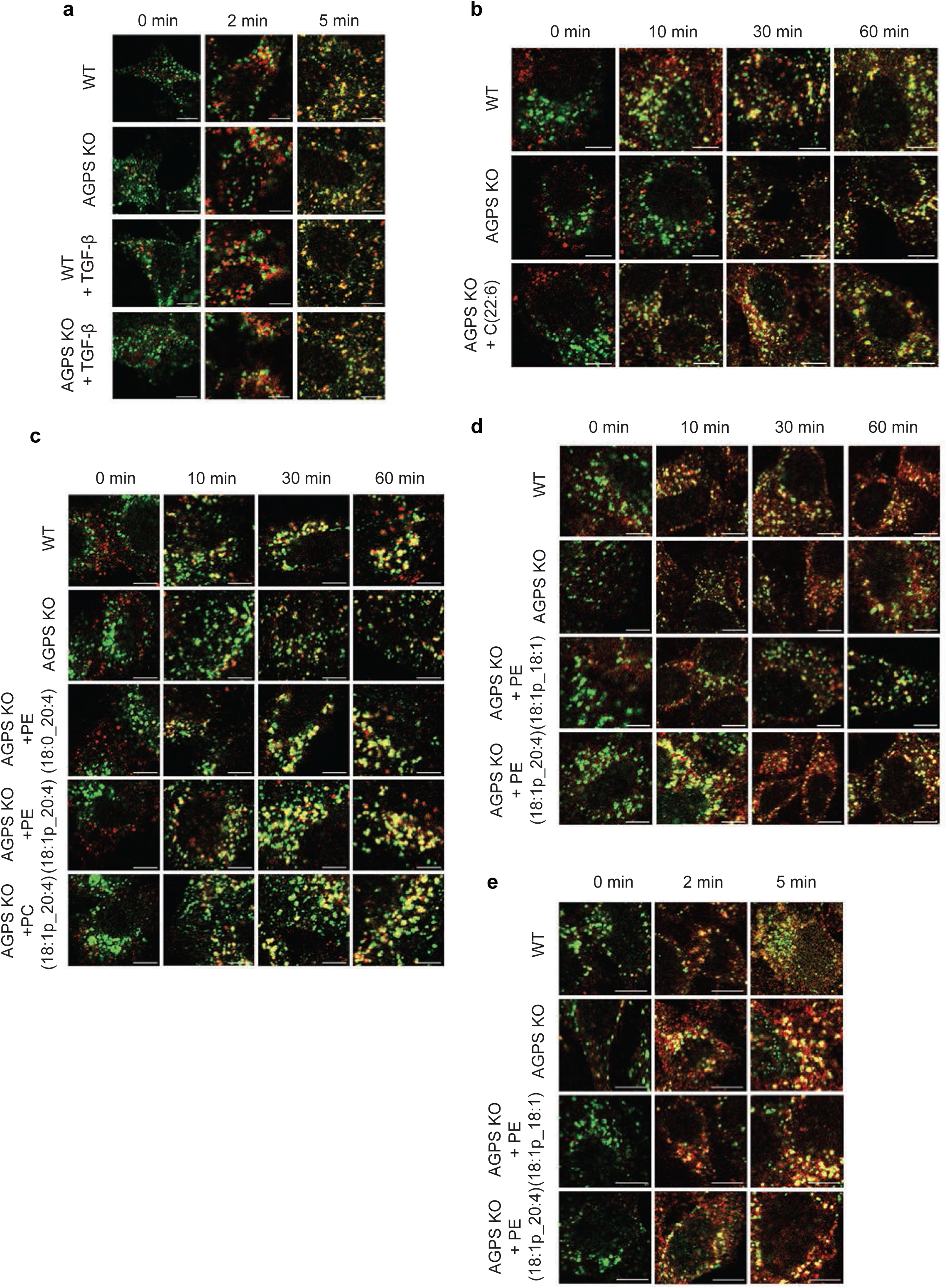
a. Representative confocal images showing EGF (red) internalization and colocalization with EEA1 (green) at 0, 2, and 5 minutes in WT and AGPS KO PyMT1099 cells +/− TGF-β pretreatment (2 ng/ml; 10 d). Scale bar: 10 μm. Cells were treated with 2 ng/ml Alexa 555-conjugated EGF. b. Representative confocal images of hyaluronate layered nanoparticle (red) and EEA1 (green) colocalization at 0, 10, 30, and 60 minutes in pB3 WT, AGPS KO, and AGPS KO supplemented with polyunsaturated fatty acid (PUFA) BSA conjugate C(22:6). Scale bar: 10 μm. c. Representative confocal images of hyaluronate layered nanoparticle (red) and EEA1 (green) colocalization at 0, 10, 30, and 60 minutes in WT, AGPS KO, and AGPS KO pB3 cells pre-treated with the indicated liposome supplementations PE (18:0_20:4), PE (18:1p_20:4), and PC (18:1p_20:4) for 16-18 hr. Scale bar: 10 μm. d. Representative confocal images of fluorescently labeled hyaluronate probe (red) and EEA1 (green) colocalization at 0, 10, 30, and 60 minutes in WT, AGPS KO, and AGPS KO pB3 cells supplemented with liposomes composed of ether PE species 18:1p_18:1 and 18:1p_20:4. Scale bar: 10 μm. e. Representative confocal images showing transferrin (red) and EEA1 (green) colocalization at 0, 2, and 5 minutes in WT, AGPS KO, and AGPS KO pB3 cells supplemented with defined ether PE species (18:1p_18:1 and 18:1p_20:4). Scale bar: 10 μm. Examined n=10 fields of cells per experimental sample for all experiments and at least two independent replicates were performed.

**Supplementary Fig. 7.**
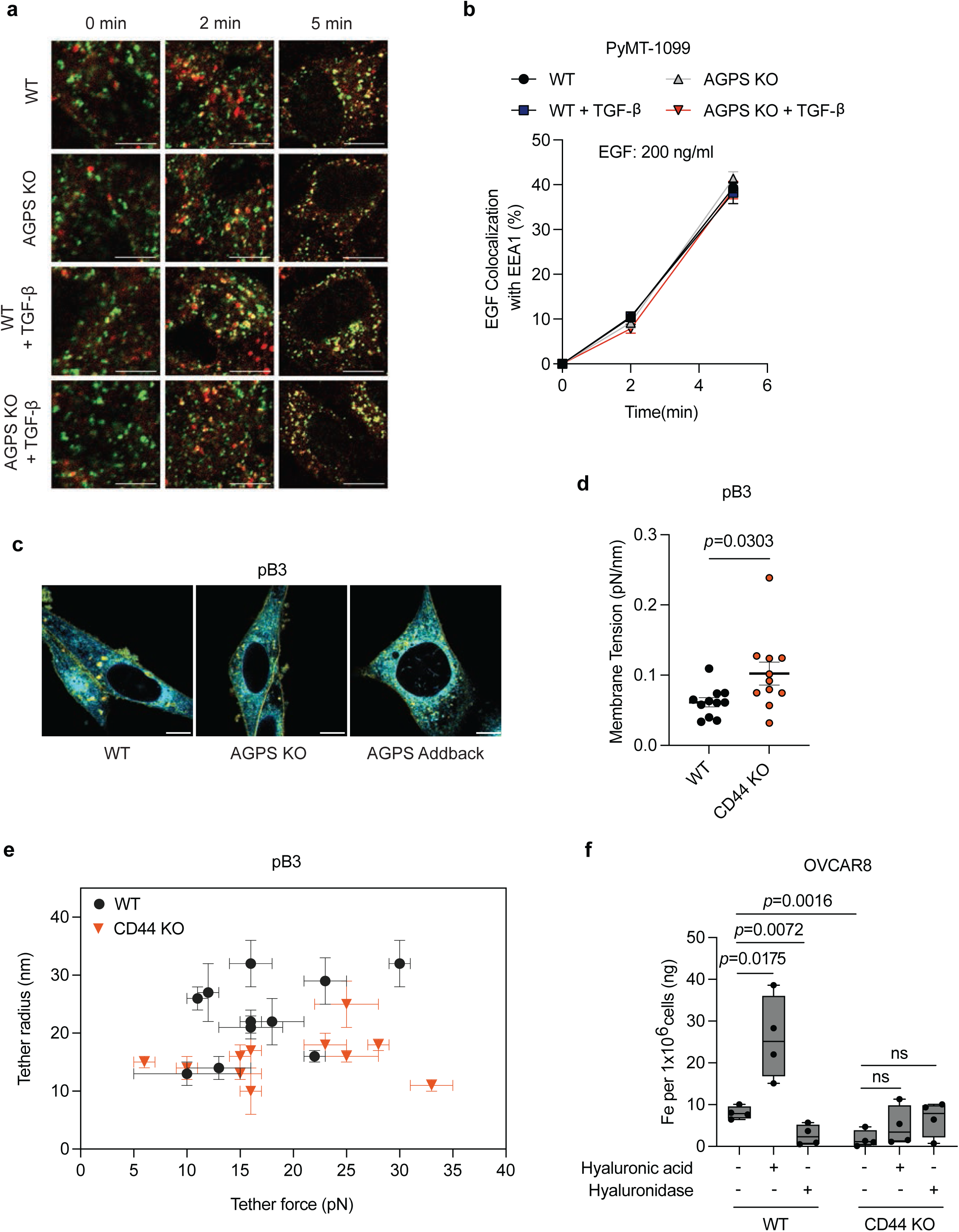
a. Representative confocal images of EGF (red) colocalization with an early endosomal marker (EEA1) (green) in PyMT-1099 WT or AGPS KO cells pretreated with 2 ng/ml TGF-β for 10 days. Cells were treated with 200 ng/ml EGF. b. Quantification of co-localization EGF as shown in panel a. All data shown as mean +/− SEM. Examined n=10 fields of cells per time point. Experiments are representative of two independent experiments. c. Representative confocal images of WT, AGPS KO, and AGPS addback pB3 cells stained with C-laurdan. The color scale represents the calculated GP ranging from 1 (blue) to 1 (red). Scale bars: 10 μm. d. Quantification of membrane tension in pB3 WT and CD44 KO. All data shown as mean +/− SEM. Statistical significance was calculated using unpaired, two-tailed ttest. e. Graph showing tether radius (R) and tether force measurements ( ) in pB3 WT and CD44 KO cells. All data shown as mean +/− SD. f. Boxplots quantifying total iron content via ICP-MS (ng Fe per 1×10 cells) in WT and CD44 KO OVCAR8 cells treated with hyaluronic acid or hyaluronidase as indicated. Data shown as mean +/− SEM (n=4). As this experiment serves as validation of a prior findings, we tested the shown comparisons using one-sided Welch’s *t*-tests without multiple-testing correction. For panel c: pB3 WT and AGPS KO cells were transduced with the respective vector control plasmids.

**Supplementary Fig. 8.**
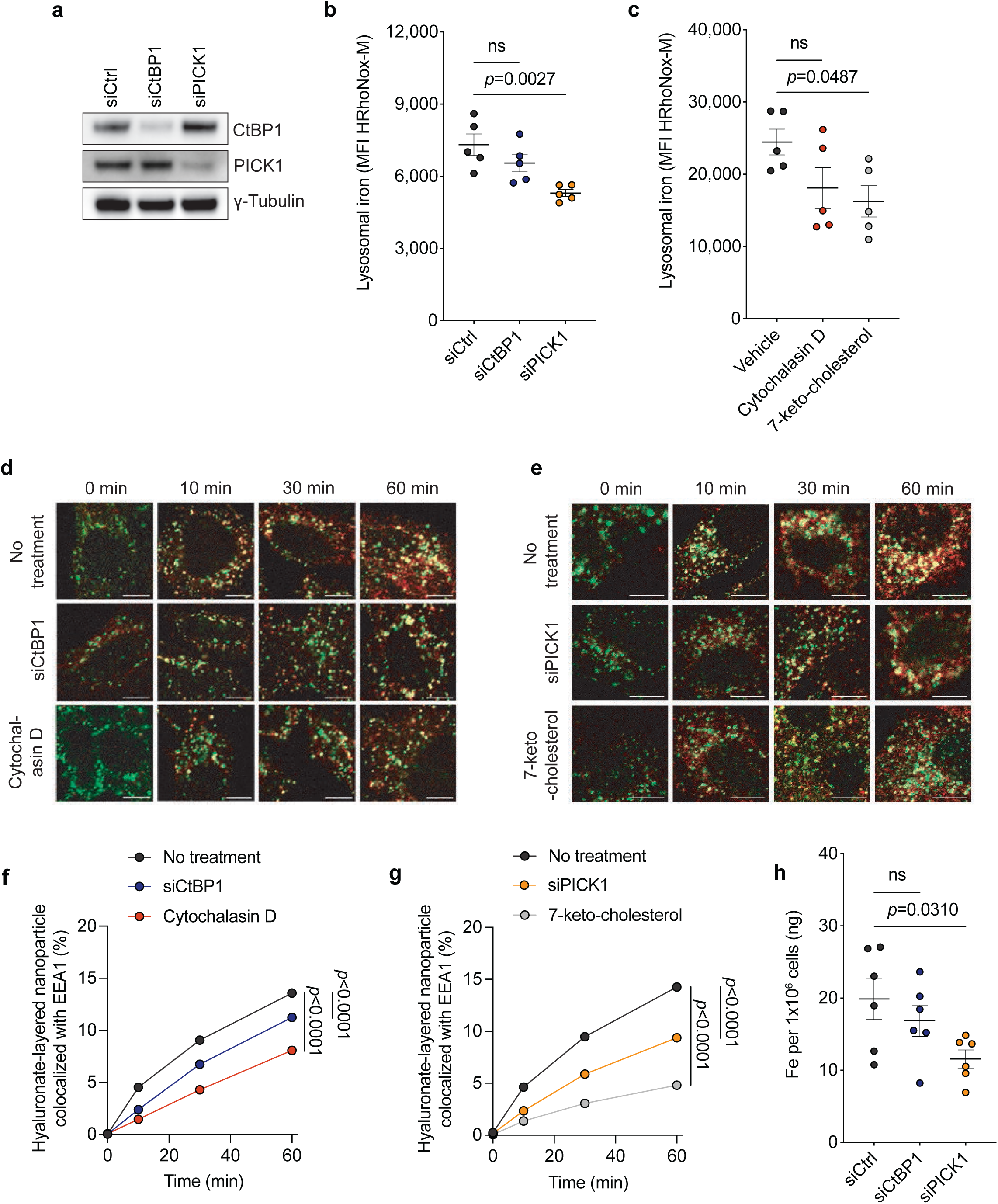
a. Immunoblot analysis of CtBP1 and PICK1 protein expression following siRNA-mediated knockdown (siCtBP1, siPICK1) or control siRNA (siCtrl) in pB3 cells. γTubulin serves as a loading control. b. Scatter plots quantifying lysosomal iron (measured as mean fluorescence intensity, MFI, of HRhoNox-M) in pB3 cells transfected with siCtrl, siCtBP1, or siPICK1 for 72 h (n=5). c. Scatter plots quantifying lysosomal iron (measured as mean fluorescence intensity, MFI, of HRhoNox-M) in pB3 cells treated with vehicle, cytochalasin D, or 7-ketocholesterol for 30 minutes (n=5). d. Representative confocal images showing hyaluronate-layered nanoparticle (red) internalization and colocalization with early endosome marker EEA1 (green) at 0, 10, 30, and 60 minutes in WT pB3 cells under the following treatments: no treatment, siCtBP1, and cytochalasin D. Data is representative of at least 2 independent experiments. Scale bar: 10 μm. e. Representative confocal images showing hyaluronate-layered nanoparticle (red) internalization and colocalization with early endosome marker EEA1 (green) in pB3 cells at 0, 10, 30, and 60 minutes in WT pB3 cells under the following treatments: no treatment, siPICK1, and 7-keto-cholesterol. Data is representative of at least 2 independent experiments. Scale bar: 10 μm. f. Quantification of the percentage of hyaluronate-layered nanoparticle labeled endosomes colocalized with EEA1 in pB3 cells over time (0–60 min) in the indicated treatment groups: No treatment, siCtBP1, or cytochalasin D. Examined n=10 fields of cells per experimental sample. Data is representative of at least 2 independent experiments. g. Quantification of the percentage of hyaluronate-layered nanoparticle labeled endosomes colocalized with EEA1 in pB3 cells over time (0–60 min) in the indicated treatment groups: No treatment, siPICK1, or 7-keto-cholesterol. Examined n=10 fields of cells per experimental sample. Data is representative of at least 2 independent experiments. h. Total iron content (ng Fe per 1×10^6^ cells) in pB3 cells transfected with siCtrl, siCtBP1, or siPICK1 as quantified by inductively coupled plasma mass spectrometry (ICM-MS) (n=6). All data shown as mean +/− SEM and statistical significance was calculated using oneway ANOVA with Dunnett’s multiple comparisons test; ns, not significant. In some cases, error bars may be obscured by data points.

**Supplementary Fig. 9.**
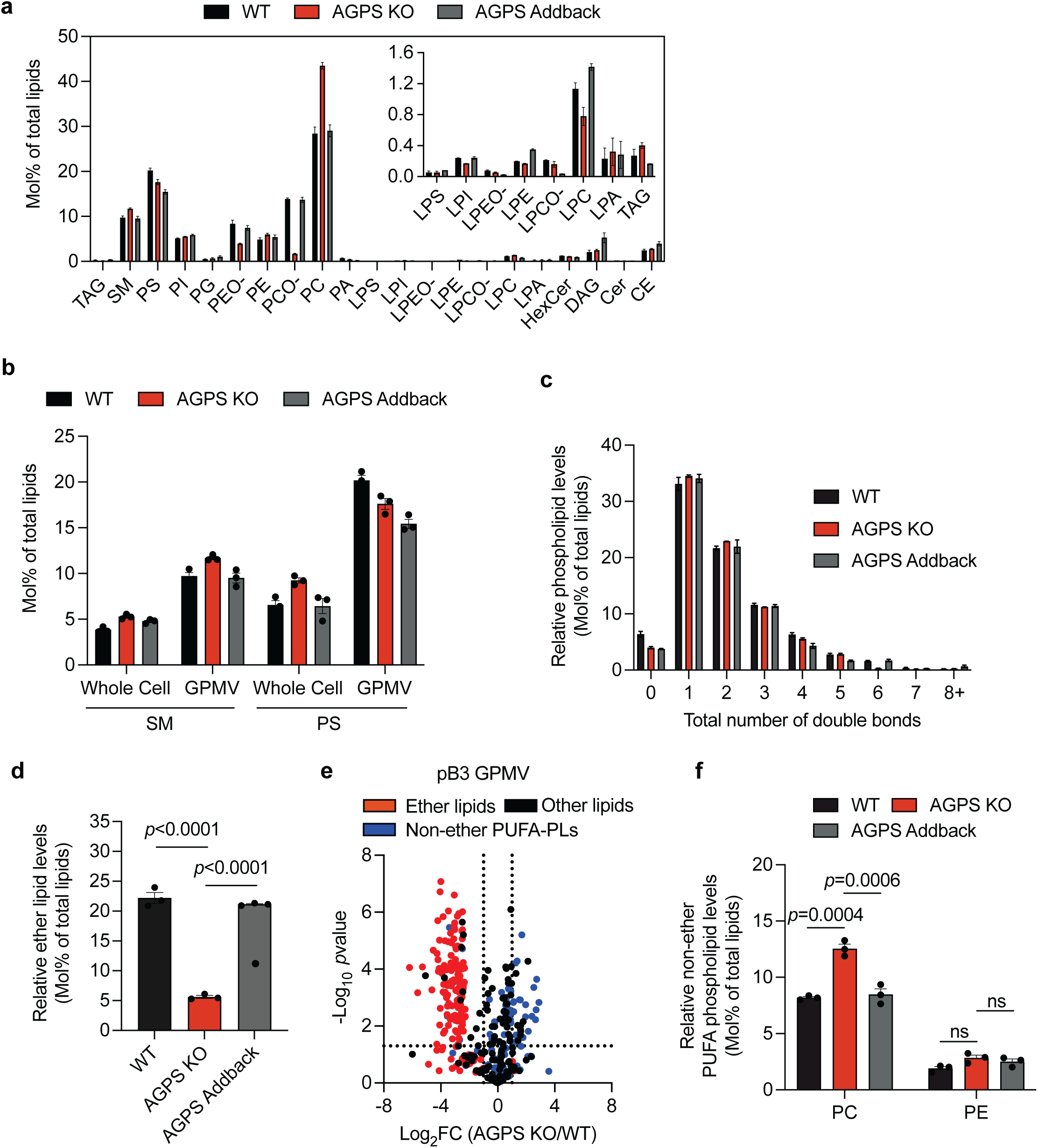
a. Bar plot showing the abundance (mol% of total lipids) of each major lipid class in isolated giant plasma membrane vesicles (GPMVs) from WT, AGPS KO and AGPS addback pB3 cells. Lipid classes include diacylglycerol (DAG), sphingomyelin (SM), phosphatidylserine (PS), phosphatidylinositol (PI), phosphatidylglycerol (PG), phosphatidylethanolamine (PE), ether phosphatidylethanolamine (PEO-), phosphatidylcholine (PC), ether phosphatidylcholine (PCO-), phosphatidic acid (PA), lysophosphatidylserine (LPS), lysophosphatidylinositol (LPI), lysophosphatidylethanolamine (LPE), lysophosphatidylcholine (LPC), ether lysophosphatidylethanolamine (LPEO-), ether lysophosphatidylcholine (LPCO-), hexosylceramide (HexCer), cardiolipin (CL), ceramide (Cer), cholesteryl ester (CE), cholesterol (Chol) and triacylglycerol (TAG). b. Enrichment in PS and SM lipid abundance (mol% of total lipids) in WT, AGPS KO, and AGPS addback pB3 derived GPMVs compared to whole cell extracts. c. Bar plot showing the distribution of relative phospholipid levels according to the total number of double bonds per lipid species in GPMVs derived from WT, AGPS KO, and AGPS addback pB3 cells. d. Quantification of total ether lipid levels (mol% of total lipids) in WT, AGPS KO, and AGPS addback pB3 derived GPMVs. e. Volcano plot showing the log_2_(fold change) in the relative abundance of various lipid species present in GPMVs upon knockout of AGPS in pB3 cells. Blue indicates non-ether linked polyunsaturated phospholipids; red indicates all ether lipids identified in lipidomic analysis and black denotes all other lipids identified. f. Quantification of the relative levels (mol% of total lipids) of non-ether polyunsaturated phosphatidylethanolamine or phosphatidylcholine lipids (PC or PE) in WT, AGPS KO, and AGPS addback pB3 cells. All data shown as mean (n=3) +/− SEM and statistical significance was calculated using one-way ANOVA with Tukey’s multiple comparisons test; ns, not significant. pB3 WT and AGPS KO cells were transduced with the respective vector control plasmids.

**Supplementary Fig. 10.**
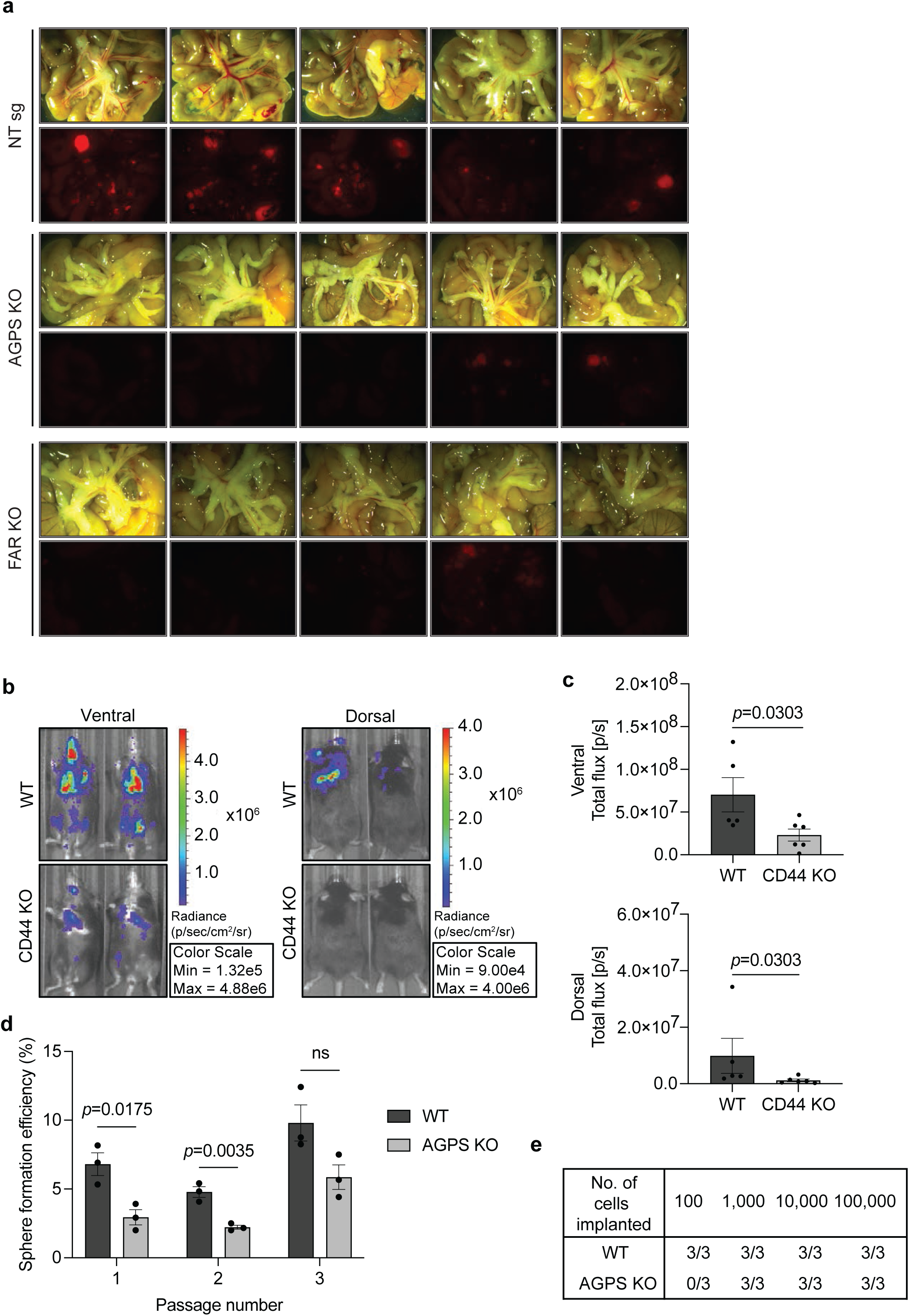
a. Bright-field (top) and fluorescence images (bottom) showing mesenteric metastases from athymic nude mice injected with tdTomato-labeled OVCAR8 NT sg, AGPS KO and FAR1 KO cells via the intraperitoneal route. b. Representative in vivo imaging system (IVIS) images of overall metastatic burden in C57BL/6 female mice following intracardiac injection of GFP-luciferized pB3 WT (n=5) and CD44 KO (n=6) cells. c. Quantification of overall metastatic burden in C57BL/6 female mice following intracardiac injection of GFP-luciferized pB3 WT (n=5) and CD44 KO (n=6) cells. Data shown as mean +/− SEM. Statistical significance was calculated using the two tailed Mann-Whitney test. d. Bar plot showing the in vitro mammosphere formation efficiency of pB3 WT and AGPS KO cells over three passages (n=3). Data is representative of three independent biological replicates. Data shown as mean +/− SEM. Statistical significance was calculated using unpaired, two-tailed t-test. e. Table showing number of pB3 WT or AGPS KO cells implanted per female C57BL/6 mouse and number of mice with tumors after 39 days.

**Supplementary Fig. 11.**
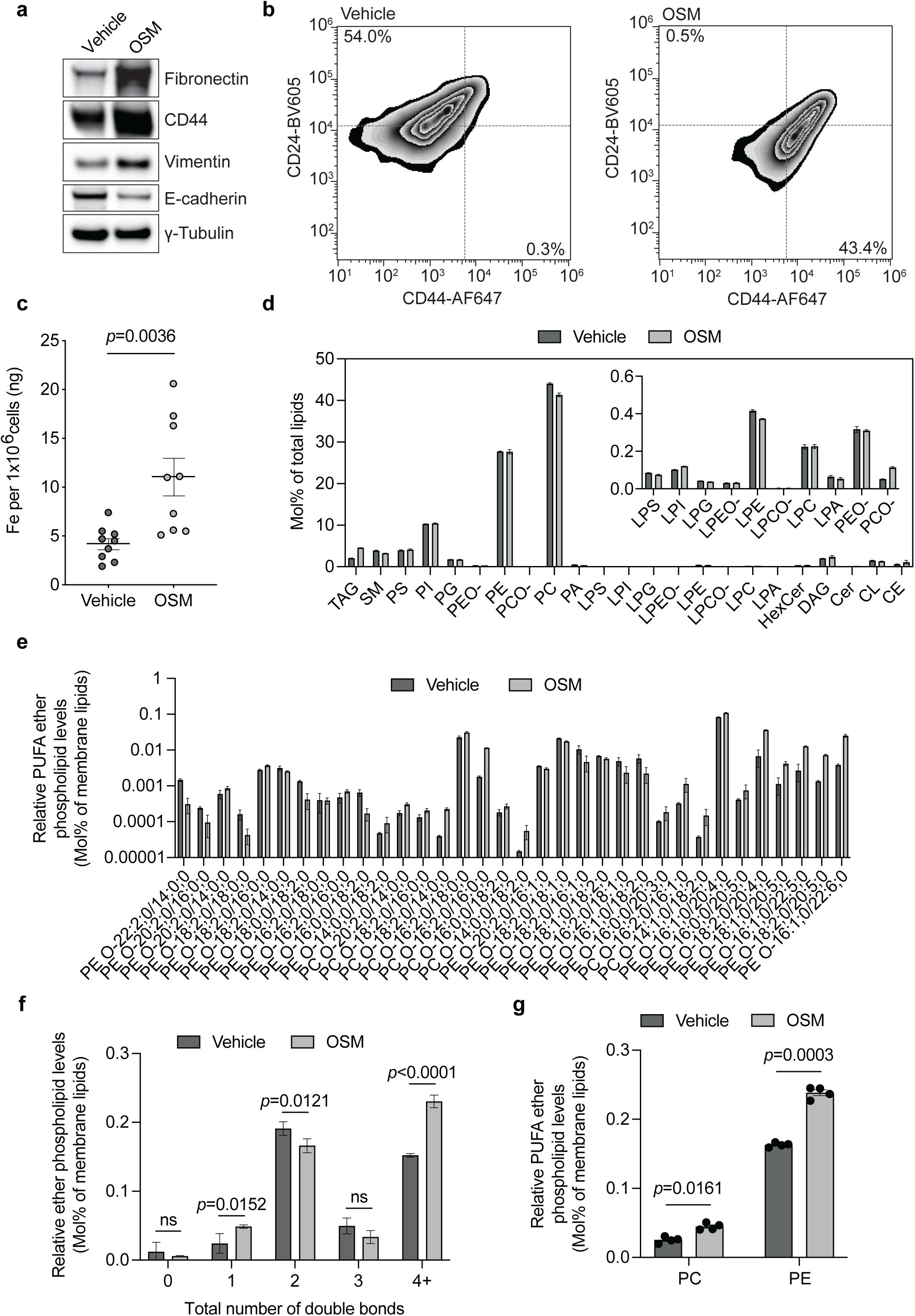
a. Immunoblot showing the expression of various EMT markers in MCF7 cells treated with vehicle control or oncostatin M (OSM, 100 ng/mL) for 72 hr. γ-Tubulin serves as a loading control. b. Representative flow cytometry plots of CD44 (AF647) and CD24 (BV605) staining in MCF7 cells treated with vehicle control or OSM (100 ng/mL) for 72 hr. Percentage of CD44^lo^/CD24^hi^ cells and CD44^hi^/CD24^lo^ is indicated. Data is representative of at least two independent experiments. c. Scatter plot quantifying total iron (ng Fe per 1×10 cells) in vehicle control or OSM (100 ng/mL; 72 hr) treated MCF7 cells (n=9). d. Bar plot showing the mol% of total lipids for each major lipid class in vehicle control and OSM treated MCF7 cells. Lipid classes include triacylglycerol (TAG), sphingomyelin (SM), phosphatidylserine (PS), phosphatidylinositol (PI), phosphatidylglycerol (PG), phosphatidylethanolamine (PE), ether phosphatidylethanolamine (PEO-), phosphatidylcholine (PC), ether phosphatidylcholine (PCO-), phosphatidic acid (PA), lysophosphatidylserine (LPS), lysophosphatidylinositol (LPI), lysophosphatidylethanolamine (LPE), lysophosphatidylglycerol (LPG), lysophosphatidylcholine (LPC), ether lysophosphatidylethanolamine (LPEO-), ether lysophosphatidylcholine (LPCO-), lysophosphatidic acid (LPA), hexosylceramide (HexCer), diacylglycerol (DAG), cardiolipin (CL), ceramide (Cer), and cholesteryl ester (CE). Inset shows low abundance lipid species. e. Relative abundance (mol% of membrane lipids) of individual ether lipid species in vehicle control and OSM treated MCF7 cells (100 ng/mL; 72 hr). f. Quantification of saturation levels (total number of double bonds) among membrane ether lipid species in vehicle and OSM treated MCF7 cells. g. Quantification of the levels (mol% of membrane lipids) of ether polyunsaturated fatty acid (PUFA) PE and PC species in vehicle control and OSM treated MCF7 cells (100 ng/mL; 72 hr). All data shown as mean +/− SEM and unless otherwise noted, statistical significance was calculated using unpaired, two-tailed t-test.

**Supplementary Fig. 12.**
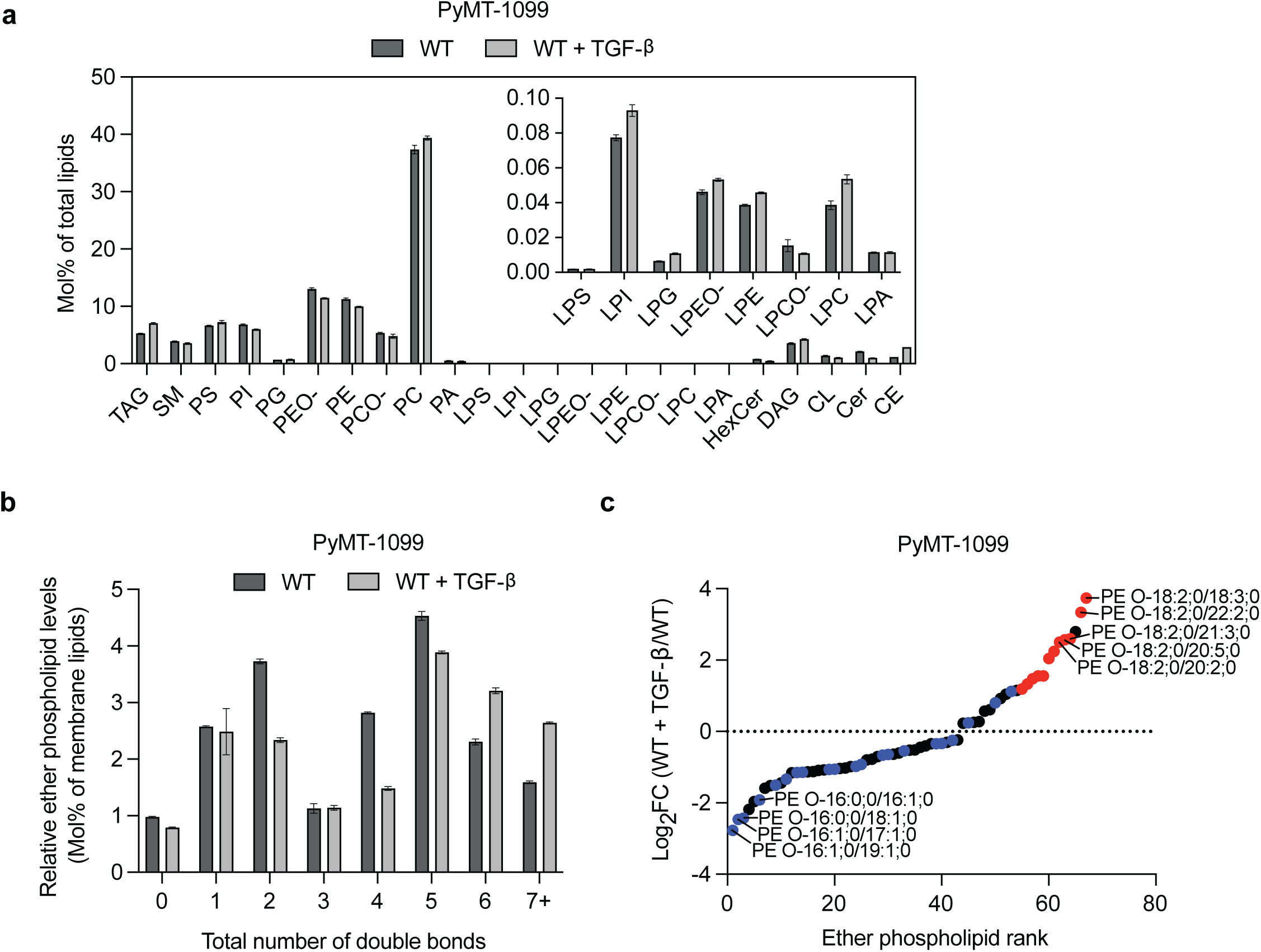
a. Relative abundance of lipid classes in TGF-β-treated (2 ng/ml; 10 d) PyMT-1099 WT cells compared to untreated control. Lipid classes include: triacylglycerol (TAG), sphingomyelin (SM), phosphatidylserine (PS), phosphatidylinositol (PI), phosphatidylglycerol (PG), phosphatidylethanolamine (PE), ether phosphatidylethanolamine (PE O-), phosphatidylcholine (PC), ether phosphatidylcholine (PC O-), phosphatidic acid (PA), lysophosphatidylserine (LPS), lysophosphatidylinositol (LPI), lysophosphatidylethanolamine (LPE), lysophosphatidylcholine (LPC), ether lysophosphatidylethanolamine (LPEO-), ether lysophosphatidylcholine (LPCO-), lysophosphatidylglycerol (LPG), lysophosphatidic acid (LPA), hexosylceramide (HexCer), diacylglycerol (DAG), cardiolipin (CL), ceramide (Cer), and cholesteryl ester (CE). Data shown as mean +/− SEM (n=3). Inset shows low abundance lipid species. b. Bar plot showing the distribution of ether phospholipids +/− SEM according to the number of double bonds per species (mol% of membrane lipids) in WT and TGFβtreated WT cells. Data shown as mean +/− SEM (n=3). c. Dot plot showing the calculated Log_2_(fold change) of ether phospholipids with an adjusted p value less than or equal to 0.02 in PyMT-1099 TGF-β-treated (2 ng/ml; 10 d) cells relative to untreated (Ctrl) cells. Red dots indicate ether phospholipids containing polyunsaturated fatty acids (PUFA) at both sn-1 and sn-2 positions. Blue dots indicate ether phospholipids containing only saturated (SA) or monounsaturated (MUFA) fatty acids at both sn-1 and sn-2 positions. Black dots indicate all other ether phospholipids.

